# Single-cell transcriptomics reveals transcriptional diversity of sea cucumber perivisceral fluid coelomocytes

**DOI:** 10.64898/2026.02.20.704403

**Authors:** Noé Wambreuse, Arnaud Lavergne, Laurence Fievez, Fabrice Bureau, Libin Zhang, Beini Deng, Guillaume Caulier, Igor Eeckhaut, Jérôme Delroisse

## Abstract

Echinoderms possess a complex immune system, primarily relying on coelomocytes – immune cells circulating in coelomic fluids. Over the last few decades, various coelomocytes have been described based on morphological features, with holothuroids exhibiting the highest diversity of cell morphotypes among the different echinoderm classes. However, while the overall immune function of these cells is broadly accepted, their respective functions remain unclear, and molecular data specific to the different cell types are still limited in the literature. In this study, we address this gap in functional information and molecular data by using single-cell RNA sequencing (scRNA-seq) on coelomocytes from the perivisceral fluid of *Holothuria forskali*. We identified 10 distinct clusters, each assumed to correspond to a distinct transcriptional coelomocyte population. Among these, cluster 0 occupies a central position relative to the others, suggesting it may represent “progenitor cells”, whereas cluster 6 is markedly divergent from all other clusters. Functional enrichment analyses revealed that some clusters ensure key immune functions, including pathogen recognition, phagocytosis, complement activation and redox balance regulation. In addition, examination of the processed samples under a microscope confirms the presence of a small proportion of recently discovered carotenocytes (7.0%) in the perivisceral fluid, a cell type rich in carotenoids. By using transcriptomics data previously obtained for this cell type by bulk RNA sequencing (bRNA-seq), it was possible to confidently identify cluster 6 as carotenocytes and provide further insights into their gene expression. While further analyses are needed to link other clusters to the different morphotypes previously described in the literature, this pioneer study presents preliminary data on the functional diversity of holothuroid coelomocytes, which could be of broad interest for a better understanding of holothuroid immunity as well as for the study of immune cell lineage evolution across deuterostomes.

## 1. Introduction

Functional characterisation of immune cell types and identification of specific marker genes for each of them have been important challenges that laid the basis of modern immunology. Indeed, nowadays it is well-documented in vertebrates that many different immune cell types exist, each with specialised functions, and that their interactions are essential to ensure the effective defence against diverse stresses (Batista and Dustin, 2013). Echinoderms are marine deuterostomes that possess an innate immune system, which is considered highly complex (Smith et al., 2018). Notably, their genomes encode hundreds of pathogen recognition receptors (PRRs), whereas vertebrate genomes generally encode only dozens of these (Hibino et al., 2016; Ward and Rosenthal, 2014). This striking diversity is thought to confer them the ability to respond specifically to a large spectrum of pathogens, despite the absence of a lymphocyte-based adaptive immune system (Ward and Rosenthal, 2014). Considering the particular phylogenetic position of echinoderms as non-chordate deuterostomes, these organisms represent an ideal system for exploring the evolution of innate immunity across the deuterostome lineage.

In echinoderms, single-cell RNA sequencing (scRNA-seq) has been primarily employed to characterise cell type differentiation in early developmental stages (*i.e.*, embryonic or larval stages), especially in sea urchins and sea stars (Massri et al., 2021; Paganos et al., 2021; Foster et al., 2022; Meyer et al., 2023). These analyses have led to the characterisation of a high number of cell subsets, including some that were attributed to “immune cells” based on immune gene expression (*e.g.,* Paganos et al., 2021; Meyer et al., 2023). However, gene expression heterogeneity within these clusters has not been investigated yet. Very recently, a first study applying scRNA-seq on coelomocytes of adult sea urchins *S. purpuratus* identified four super-clusters which would correspond to the four main coelomocyte morphotypes in echinoids (Kell et al., 2026). However, this study primarily aims to understand the response of these different clusters to UVB-induced stress, without investigating the respective gene expression among cell types, and the cluster identification remains to be validated. Besides, scRNA-seq data on adult echinoderms are still sparse in the literature: in holothuroids, scRNA-seq was employed to characterise different pigmented cells in the integuments of *Apostichopus japonicus*, as well as different neuronal cell populations around the circumoral nerve ring (Xing et al., 2023; Zheng et al., 2024). In addition, cell differentiation from the coelomic epithelia during intestine regeneration was studied in *Holothuria glaberrima*, emphasising several clusters corresponding to coelomocytes, notably characterised by the expression of fibrinogen-related proteins (Medina-Feliciano et al., 2025). However, the link between different morphotype expression and clusters identified as coelomocytes was not investigated in this study. Although all these studies have truly revolutionised the study of cell lineage in echinoderms, the characterisation of coelomocytes using this valuable tool has yet to be achieved.

Among echinoderms, it appears that holothuroids show the greatest diversity of coelomocyte morphotypes, with generally six accepted cell types (Hetzel 1963; Smith et al., 2018), although less restrictive classifications report up to nine morphotypes (Queiroz and Custódio, 2024 for review). These six cell types include phagocytes, spherule cells, progenitor cells, haemocytes, fusiform cells and crystal cells. However, while these different cell types were extensively studied morphologically (*e.g.,* Endean, 1958; Xing et al., 2008; Caulier et al., 2020; Queiroz et al., 2022; Wambreuse et al., 2025a, b), very little information exists on their respective function, and transcriptomics data associated with the different cell types are still very limited. Recently, several attempts have been made to provide such transcriptomics data on a subset of coelomocytes using bulk RNA sequencing (bRNA-seq) on enriched populations of different types of coelomocytes in holothuroids (Yu et al., 2023; Wambreuse et al., 2025a). Yu et al. (2023) compared two subsets of coelomocytes, namely “spherical cells” and “lymphocyte-like cells” in *A. japonicus*, showing that spherical cells overexpressed genes involved in phagocytosis and humoral response in comparison to lymphocyte-like cells. In addition, comparing coelomocyte gene expression between the two main body fluids of *Holothuria forskali*, namely the hydrovascular and perivisceral fluids, has recently enabled us to gain knowledge on the carotenocytes (Wambreuse et al., 2025a). This cell type, containing carotenoids, is highly enriched within the hydrovascular fluid but present in very low proportions in the perivisceral fluid. Although such studies provide valuable data, they constitute an approximation of the gene expression at the cell-type resolution, which can be surpassed by using scRNA-seq. In this study, we employed, for the first time, scRNA-seq on perivisceral coelomocytes of a holothuroid (*H. forskali*) to assess the transcriptional landscape of coelomocyte types and their respective immune function. Although the identification of some clusters still requires validation to link them with coelomocyte morphotypes, it provides valuable data on the coelomocyte functional diversity and will constitute a first step in defining functional coelomocyte types in holothuroids.

## 2. Materials and Methods

### 2.1. Maintenance of organisms

Specimens of *Holothuria forskali* Delle Chiaje, 1823 were obtained from the Biological Station of Roscoff, which collected them by SCUBA diving on the shallow rocky seafloor of Morlaix Bay (Brittany, France). Once delivered to the University of Mons (Belgium), specimens were maintained in a 400 L tank in closed circuit with seawater at 33-35 psu and 15°C with a circadian cycle reproduced by UV-enhanced fluorescent tubes from 8 am to 8 pm. Before being processed, the specimens were acclimatised for at least two weeks without any handling. During this time, sea cucumbers were fed once a week with a mix of dried algae in agar-agar-based gel.

### 2.2. Coelomocyte collection and preparation

Coelomocytes from the perivisceral fluid of *H. forskali* were collected as described in Wambreuse et al. (2025b). Briefly, an incision was made on the *bivium* (*i.e.,* dorsal side) of the specimen between two radial canals, and the perivisceral coelomic fluid was harvested in a 15 ml tube. This tube was kept on ice to prevent coelomocytes from clotting. Importantly, only female individuals were considered, sexed based on the aspect of their gonads (as per Tuwo and Conand, 1992), to avoid any contaminating spermatozoa, as these are commonly accidentally captured when harvesting coelomocytes in male individuals (Caulier et al., 2024). The sample was then filtered on a 70 µm-pore Flowmi filter to discard cell aggregates. In parallel, 20 µl were pipetted to carry out a cell count on a hemacytometer and estimate the number of cells in the tubes, allowing the resuspension of the cell pellet in a proper volume to obtain a suitable concentration for single-cell library preparation, namely 10^6^ cells per ml. The sample was then centrifuged, slower than usual, at 350 × g and 4°C to avoid cell stress. The cell pellet was resuspended in calcium and magnesium-free Phosphate Buffered Saline (PBS; pH = 7.4) + 0.04% of Bovine Serum Albumin (BSA). The sample was filtered a second time through a 40 µm-pore size Flowmi filter, and cell viability was assessed using trypan blue colouration on a CellDrop reader. Note that all these steps were completed as quickly as possible (approximately 30 minutes) to minimise cell degradation.

To estimate the proportion of the different cell types in the processed sample, a cell count was carried out on a hemacytometer with a remaining volume of the same cell suspension processed through the Chromium instrument. Cell types were identified visually under a microscope based on a previous study of coelomocyte morphotypes in this species (Wambreuse et al., 2025a). This proportion was compared to the previous results of cell count obtained from six individuals. Note that while the cell count was carried out in the same way, for the former ones, calcium- and magnesium-free artificial seawater containing EDTA (CMFSW + EDTA: 460 mM NaCl; 10.7 mM KCl; 7 mM Na_2_SO_4_; 2.4 mM NaHCO_3_; 20 mM HEPES; 70 mM EDTA; pH = 7.4) was added to avoid cell aggregation (Smith et al., 2019). Note that it was not possible to use EDTA for the sample processed for the scRNA-seq, as EDTA is an inhibitor of enzymatic reactions needed for library preparation.

### 2.3. Cell separation in GEMs, library preparation and sequencing

Once one sample met the concentration and cell viability criteria, it was processed directly on the 10x Chromium instrument. Briefly, cells were separated in a microfluidic system and confined within Gel Beads in Emulsions (GEMs) that contain all necessary reagents and enzymes to generate a cDNA library for each cell. These cDNAs will contain a barcode, unique to each initial bead (thus unique to each cell), as well as a unique molecular identifier (UMI) unique to each mRNA. These short sequences are essential in the scRNA-seq technology as they enable matching transcripts to the cells (via the barcodes) and reduce amplification redundancy (via the UMIs) in post-analysis. After GEM dissociation, cDNA amplification, purification, and preparation for sequencing, the cDNA library was sequenced using an Illumina NovaSEQ 6000 platform.

### 2.4. Mapping of reads on the *de novo* transcriptome and generation of matrix counts

Raw FASTQ files generated from a droplet-based single-cell platform (10x Genomics Chromium) were used as input. scRNA-seq data were processed using Alevin, a tool within the Salmon v1.10.3 suite (Srivastava et al., 2019), designed for efficient and accurate quantification of barcoded single-cell transcriptomes.

In the absence of a reference genome for *H. forskali*, sequenced reads were mapped to a high-quality *de novo* transcriptome previously generated from coelomocytes of the same species (Wambreuse and Bossiroy, 2025; https://doi.org/10.6084/m9.figshare.c.7839038). This transcriptome consists of a set of “predicted genes” called unigenes, which may include transcript fragments, isoforms, or redundant sequences. For clarity and consistency, these features are referred to as “genes” throughout this manuscript. However, caution is warranted when interpreting the number of marker genes identified for specific gene families, as redundancy within the transcriptome could lead to inflated gene counts and duplicated marker genes.

### 2.5. Generation of clusters and identification of marker genes

scRNA-seq data were processed and analysed using R (v4.4.2) and the Seurat package (v5.3.0) for downstream analysis (Hao et al., 2024). Raw count matrices were imported from Alevin output files to create a Seurat object with filtering parameters set to retain cells expressing at least 100 genes and genes detected in a minimum of 3 cells.

#### 2.5.1. Quality control and scaling

Basic quality control metrics were assessed, including the number of RNA molecules (nCount_RNA) per cell and the number of detected genes (nFeature_RNA). The data were normalised with a default scale factor of 10,000. The 2,000 most variable features were identified with “FindVariableFeatures” and used for scaling, while regressing out the total RNA count per cell.

#### 2.5.2. Dimensionality reduction and clustering

Principal component analysis (PCA) was performed on the scaled data, and the first 30 principal components were used to construct a Shared Nearest Neighbour (SNN) graph via “FindNeighbors”. Clusters were identified using “FindClusters” with a resolution of 0.25 (a value which yields a reasonable number of clusters, consistent with the number of described coelomocyte morphotypes: Queiroz and Custódio, 2024). Uniform Manifold Approximation and Projection (UMAP) was applied for dimensionality reduction and visualisation, using the same principal components and a minimum distance of 0.35.

#### 2.5.4. Detection of mitochondrial expression

As no list of mitochondrial genes is directly linked to the *de novo* transcriptome, detection of mitochondrial genes in the transcriptome was done based on general annotation previously obtained for the transcriptome (see Wambreuse et al., 2025a). To do this, genes present in the mitochondrial genome of *H. forskali* (Perseke et al., 2010) were searched among the transcriptome, including genes coding for “NADH dehydrogenase”, “cytochrome B”, “cytochrome c oxidase” and “ATP synthase”. Only genes with an e-value < 10^-20^ were retained to ensure a reliable annotation. The list of mitochondrial genes was then loaded on R, and the expression percentage of mitochondrial genes (based on gene counts) in each cell was calculated using the function “PercentageFeatureSet”. These percentages were finally mapped on the UMAP, and the distribution of these percentages was represented among the cells. For comparisons, cells having a mitochondrial gene expression proportion < 20% were retained, reclustered, and re-embedded using UMAP. However, we decided to retain the unfiltered dataset for downstream analysis due to our low number of cells for some clusters and to avoid artificially added bias.

#### 2.5.5. Doublet detection

The distribution of putative cell doublets was assessed as part of the quality control using the DoubletFinder package (v2.0.6) in R (McGinnis et al., 2019). Briefly, doublet detection was performed on the Seurat object after normalisation, dimensionality reduction, and clustering. The first 20 principal components (PCs) were used to define cell–cell similarity. To determine the optimal neighbourhood size parameter (pK), a parameter sweep was conducted using the paramSweep and summarizeSweep functions, and the pK value maximising the BC metric was selected. Artificial doublets were generated at a proportion of 25% relative to the number of real cells (pN = 0.25), as recommended by McGinnis et al. (2019). The expected number of real doublets (nExp) was estimated as 10% of the total number of cells, which is consistent with the upper range of what is typically encountered with 10x technology (McGinnis et al., 2019). DoubletFinder was then applied to compute a per-cell doublet score (pANN), representing the proportion of artificial nearest neighbours, and to classify cells as either singlets or doublets. Doublet classifications and pANN scores were visualised on the UMAP to assess their spatial distribution across clusters. In addition, the proportion of predicted doublets and the mean pANN score were quantified for each cluster to identify clusters potentially enriched in doublet-like transcriptional profiles. For comparisons, cells classified as singlets were retained, reclustered, and re-embedded using UMAP. However, we decided to retain the unfiltered dataset for downstream analysis due to our low number of cells for some clusters and to avoid artificially added bias.

#### 2.5.6. Marker gene identification and visualisation

Cluster-specific marker genes were identified using the “FindAllMarkers” function with the following thresholds: adjusted p-value (Padj) < 0.01 and average log_2_(fold-change) (avgFC) ≥ 0.5. The top five markers per cluster were selected based on avgFC, and dot plots were generated to visualise their expression using a custom plotting function. The Nr annotation of each marker gene was added from an external gene annotation file obtained from Wambreuse et al. (2025a). As many marker genes were not annotated, the marker gene files were filtered to keep only marker genes with an annotation (*i.e.,* filtering of “NA” as annotation). Gene expression signatures were constructed based on the five top marker genes per cluster. For each cluster, a score was computed using “PercentageFeatureSet” to evaluate the proportion of expressed genes in the respective signature. This signature was then represented on UMAP using a colour scale to assess the degree of specificity of marker genes for each cluster.

### 2.6. Functional enrichment analysis

To functionally characterise the transcriptional profiles of each cell cluster, we performed a series of enrichment analyses using curated gene annotations from KEGG (Kyoto Encyclopedia of Genes and Genomes) and Gene Ontology (GO) databases, encompassing the three main GO categories: Biological Process (BP), Cellular Component (CC), and Molecular Function (MF), previously obtain for the *de novo* transcriptome (Wambreuse et al., 2025a).

Cluster-specific gene lists were extracted and formatted into named vectors or lists. Enrichment analyses were conducted using the “compareCluster” or “enricher” functions from the clusterProfiler package (Yu et al., 2012), which supports both GO and KEGG over-representation analysis. GO annotations were parsed and filtered using ontology information retrieved via the GO.db package (Carlson, 2017). For custom enrichment, term–name (TERM2NAME) mappings were constructed directly from the annotation tables. For the KEGG enrichment, KEGG Orthology (KO) identifiers were used, and enrichment was carried out using the “KO” organism reference within KEGG, enabling a broad taxonomic coverage. For the GO enrichment, only annotated terms corresponding to the desired ontology category (BP, CC, or MF) were retained using the “Ontology” function. Gene-term associations were then used to perform enrichment analysis for each cluster.

The resulting enrichment outputs were compiled in **Table S4**. For visualisation, the top five enriched terms per cluster (based on Fold Enrichment value) were selected and plotted using ggplot2, with dot plots displaying term names, number of associated genes (point size), and adjusted significance (-log_10_(q-value)) as a colour gradient. Note that for this visualisation, no filtering was initially applied on q-values (cutoffs set to 1) to retain at least one term or pathway for each cluster. However, significantly enriched pathways are shown in **Table S4** (in green) with a cutoff set as q-value < 0.05.

### 2.7. Identification of carotenocytes clusters based on transcriptomic datasets

In order to identify a cell cluster corresponding to carotenocytes, expression of single-cell datasets was compared with a differential expression analysis acquired from bRNA-seq between the hydrovascular fluid, enriched in carotenocytes and perivisceral fluid, in which carotenocytes are almost absent (Wambreuse et al., 2025a). To avoid misunderstanding related to the cellular composition of these body fluids, these two samples will be referred to as “CArotenocyte-Enriched sample – CAE” and “Other Coelomocyte-Enriched sample – OCE”, respectively. This dataset comprises a list of differentially expressed genes (DEGs) and their corresponding expression values in the two cell populations. To have a first idea of a candidate cluster for carotenocytes, we check the number and the proportion of marker genes of each cluster matching DEGs, as well as whether those were more expressed in OCE or CAE samples. Then, a curated list of matching genes with the best candidate cluster (the one having the highest proportion of matching DEG overexpressed CAE) was used to compute a signature score via the “PercentageFeatureSet” across these genes. The results were visualised on the UMAP, and a violin plot comparing this signature between the different clusters was generated.

To support these data, expression value (FPKM) of marker genes between OCE and CAE from the best candidate cluster was visualised on a heatmap in MetaboAnalyst V6.0 (after log transformation and autoscaling). The list of marker genes matching DEGs overexpressed in CAE was also visualised on a heatmap (in the same way as explained above), along with their Nr annotation, to provide functional information on these genes. For this heatmap, no clustering was performed as genes were ordered according to their avgFC for the cluster.

Finally, to further identify cell identities, the SingleR package was used for reference-based cell type annotation. The Seurat object was converted to a SingleCellExperiment and normalised with “logNormCounts”. Reference profiles were generated from bRNA-seq data comparing the two conditions (*i.e.*, DEG expression between OCE and CAE), and annotations were assigned using “SingleR” with these reference labels (*i.e.,* mean expression in the two conditions). The results were visualised using UMAP and bar plots showing the proportion of cells that were assigned either to CAE or OCE in each cluster.

## 3. Results and discussion

### 3.1. ScRNA-seq revealed different transcriptional coelomocyte populations

Coelomocyte classification has long been a challenging exercise, subject to much controversy (Hetzel, 1963; Chia and Xing, 1996; Caulier et al., 2024; Queiroz and Custódio, 2024; Queiroz et al., 2025; Wambreuse et al., 2025a). This difficulty is partially due to the lack of efficient techniques that separate coelomocyte morphotypes to perform molecular and functional characterisation (Smith et al., 2019; Queiroz and Custódio, 2024). scRNA-seq enables overcoming this challenge as it provides transcriptomic information at the single-cell level, without the need for prior cell purification (Bump and Lubeck, 2023). In this way, we employed for the first time this tool on circulating coelomocytes in holothuroids (**Fig. 1A**), which allows us to identify ten clusters, assumed to correspond to ten functional cell types (**Fig. 1**). However, although scRNA-seq is a powerful tool that can be applied to non-model organisms, the lack of molecular data associated with different clusters makes it difficult to link the different clusters obtained with morphotypes previously described (Bump and Lubeck, 2023).

**Fig. 1.**
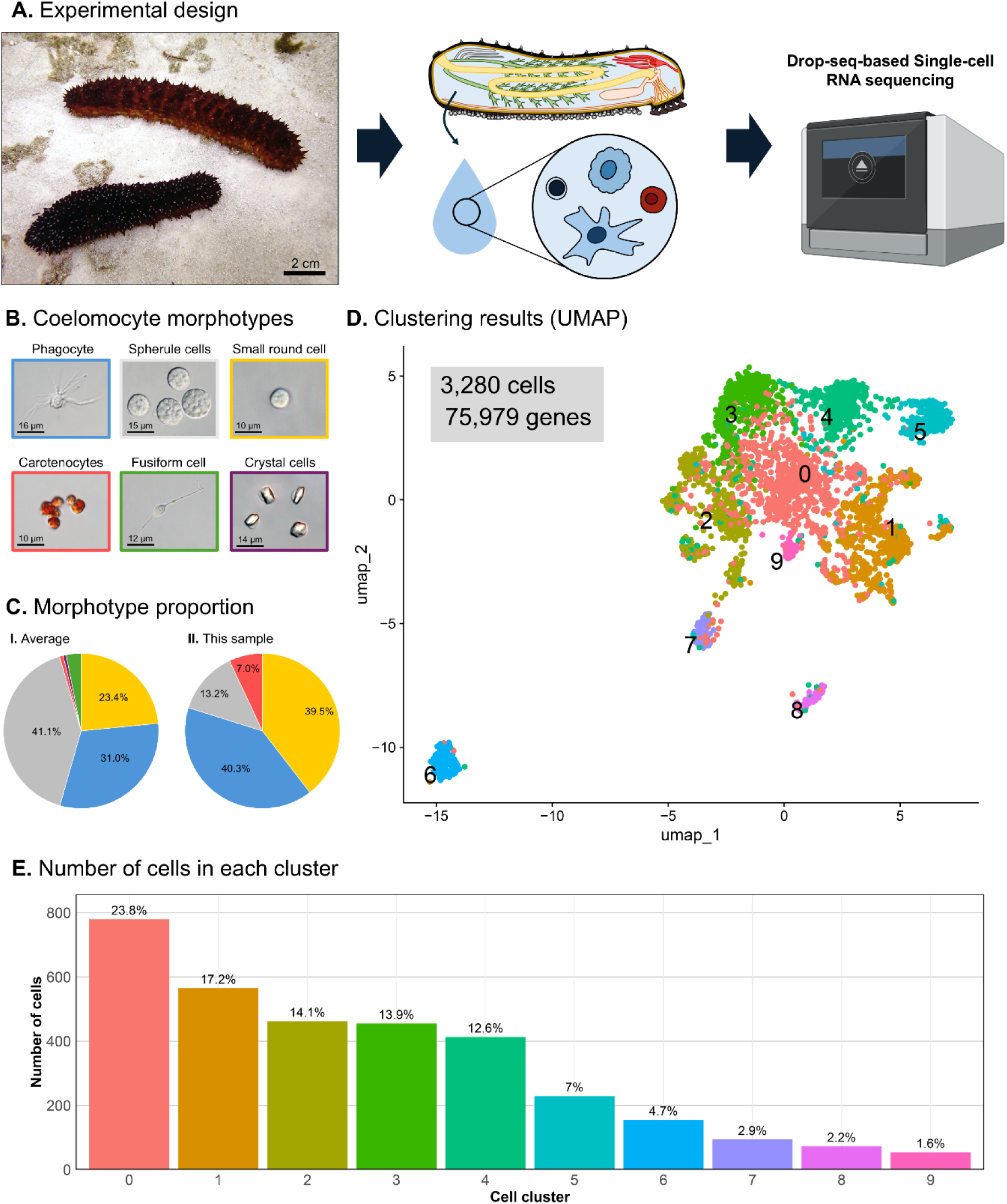
Single-cell sequencing analysis (scRNA-seq) on perivisceral coelomocytes in *Holothuria forskali*. **A.** Experimental design: coelomocytes were extracted from the perivisceral cavity of specimens of *H. forskali*, and processed as a pool of coelomocyte morphotypes for Drop-seq-based scRNA-seq using the 10x Genomics technology. **B.** Main coelomocyte morphotypes found in the coelomic fluids of *H. forskali*. **C.** Proportion of these morphotypes (I) generally observed (mean proportion; n = 6 — data from Wambreuse et al., 2025a), and (II) established in the processed sample (the colour corresponds to those surrounding cell morphotype in B). **D.** Clustering results visualised on a UMAP (Uniform Manifold Approximation and Projection), numbers and colours correspond to the different cell clusters. **E.** Number and proportion of cells in each cluster.

In *H. forskali*, the coelomic fluids (*i.e.*, perivisceral and hydrovascular fluids) are composed of six main cell morphotypes, including phagocytes, spherule cells, small round cells, carotenocytes, fusiform cells and crystal cells (**Fig. 1B**; Wambreuse et al., 2025a). Phagocytes and spherule cells are generally the predominant cell types at 31.0 ± 10.7% and 41.1% ± 9.0%, respectively (n = 6; **Fig. 1C**; data from Wambreuse et al., 2025a). Small round cells (SRCs) are also relatively abundant at 23.4 ± 7.36%, whereas other cell types account for less than 5%. Carotenocytes in particular are poorly represented in the perivisceral fluid (< 1%), but are highly abundant in the hydrovascular fluid (80.62 ± 5.14%; Wambreuse et al., 2025a). Microscopic examination of the processed sample revealed slightly different proportions: phagocytes and SRCs were the dominant types (40.3% and 39.5%, respectively), with spherule cells being less represented than usually observed (13.2%). Interestingly, a higher proportion of carotenocytes was observed than usual, reaching 7.0% of the total count. Although this number is higher than previously observed in the perivisceral fluid, similar proportions were occasionally found in this species (up to 20%; NW personal observation). No fusiform cells and crystal cells could be detected visually in this sample (**Fig. 1C**). However, it is important to consider that processing of cells for the Chromium instrument required some modifications to the protocol generally used for cell counts, which may have influenced the proportion of the different cell types in comparison to cell proportions usually encountered. Moreover, these two cell types are less common than others (< 5%) and inconsistently detected across individuals (Queiroz and Custódio, 2024; Wambreuse et al., 2025a), which may make them more difficult to capture. Note that crystal cells were recently suggested to be phagocytes that have phagocytosed biogenic crystals considered as metabolism by-products (Ma et al., 2026), hence they may not constitute a specific transcriptional population per se.

The scRNA-seq analysis resulted in 3,280 cells and 75,979 expressed “genes” (*i.e.,* predicted genes from the *de novo* transcriptome), corresponding to 45.4% of the predicted genes of the original transcriptome (containing 167,199 predicted genes). The clustering analysis revealed 10 clusters on the UMAP (**Fig. 1D**), showing a super cluster composed of clusters 0, 1, 2, 3, 4, 5 and 7, with clusters 8 and particularly cluster 6 being isolated from all other clusters. Interestingly, the supercluster is organised around cluster 0, corresponding to the most abundant cluster (**Fig. 1D**). The percentage of cells in each cluster ranges greatly from 1.6% (cluster 9) to 23.8% (cluster 0) (**Fig. 1E**). This particular topography around cluster 0 suggests that this cluster could correspond to undifferentiated coelomocytes. This characteristic is often attributed to SRCs, also referred to as progenitor cells, due to their undifferentiated morphology (Chia and Xing, 1996; Eliseikina and Magarlamov, 2002; Caulier et al., 2020). The fact that cluster 0 could correspond to this cell morphotype is also supported by the cell count, which also shows SRCs as the most abundant cell type. However, these proportions should be interpreted carefully, as it is known that different cell types do not have the same capture efficiency in Drop-seq-based scRNA-seq (*e.g.*, Borrelli et al., 2024). Mean gene expression and percentage of cells expressing each gene in each cluster are shown in **Table S1**.

Quality control analysis showed the distribution of RNA molecules (UMIs) and detected genes (features) per cell (**Fig. S1**). Dotted lines indicate thresholds of 300 UMIs and 300 features per cell, with most cells falling well above these cutoffs, reflecting the overall good quality of the dataset (**Fig. S1A** and **B**). In addition, a strong positive correlation between the number of UMIs and the number of detected features per cell is observed (**Fig. S1C**), further supporting data quality. Finally, **Fig. S1D** shows PCA based on the normalised expression data, where the first principal component (PC1) clearly separates cells from cluster 2, and the second component (PC2) distinguishes cluster 6 from the rest, indicating biologically meaningful variance across cell populations.

As an additional quality metric, the proportion of mitochondrial gene expression in the datasets was calculated. The mean mitochondrial gene expression percentage was 11.2 ± 9.6 % (**Fig. S2**). This percentage was not equally distributed between clusters, with clusters 0, 8 and 3 having the highest mean mitochondrial gene expression proportion, at 15.0 ± 12.2% and, 13.9 ± 10.6% and 13.2 ± 8.6%, respectively (**Fig. S2A-B**). In addition, we used the threshold of a mitochondrial gene expression proportion > 20% as a possible filtering parameter, which constitutes an upper range of value that can be found in cell types having a high mitochondrial expression in mammals with normal homeostasis (Osorio and Cai, 2021). By including all clusters, 85.1% of cells have a proportion < 20%. Again, this value was highly variable between clusters, with cluster 0 having the highest, namely 27.9% (**Fig. S2B-C**). Although these proportions could seem high in comparison to mammalian immune cell standard (generally around 5%), it should be highlighted that mitochondrial gene expression is highly species-and cell-type-dependent and that traditional QC cannot systematically be applied to non-model organisms (Osorio and Cai, 2021; Bump et al., 2023). Therefore, it is unknown whether this high proportion is related to the stress, particularly for that population, or if it constitutes a specific feature for this population.

Finally, DoubletFinder was used to assess the potential of each cluster to constitute a mix of several coelomocyte types (**Fig. S3**). pANN score was used to classify the cell as a singlet or a doublet – this score ranges from 0 to 1, with a higher value meaning a higher chance of being a doublet. The expected number of doublets was artificially fixed at 10%, meaning that the 10% of the cells with the highest pANN score, namely 380 cells in our dataset, were classified as doublets. Representation of pANN score on the UMAP revealed that it is not uniform across clusters, with clusters 0 and 9 having a low pANN score (∼ 0.1) and some cells from clusters 2, 3, 7 and 8 having a higher score (> 0.35; **Fig. S3C and D**). When looking at the proportion of doublets detected per cluster, the four same clusters have a higher proportion than the others, namely 39.3% for cluster 2, 22.2% for cluster 3, 19.1% for cluster 7 and 9.6% for cluster 8. Other clusters display lower proportions, under 2% (**Fig. S3E and F**).

While these different quality controls could reveal some possible filtering to apply to the dataset, we have decided not to apply a filter based on doublet identification and expression of mitochondrial genes. This was done in order to keep as much biological information as possible and to avoid artificially biased filtering for cells, for which information on gene expression is very limited. Furthermore, filtering can greatly impact the number of cells, which can also weaken marker gene identification and functional enrichment analyses. However, **Fig. S4** shows the effect of applying these filters, each or combined, on the UMAP and clustering. Filtering according to a mitochondrion gene expression > 20% result in a division of cluster 1 into two clusters and a total of 2,791 cells (**Fig. S4B**). Filtering according to the 10% of doublets results, in contrast, in the fusion of clusters 2 and 3, yielding a total of 2,952 (**Fig. S4C**). Finally, combining these two filters has the same effect as the mitochondrial expression filter alone, namely the division of cluster 1 into two clusters, with a total of 2,512 genes (**Fig. S4C**). Despite these modifications, it can be observed that the general configuration of the UMAP is conserved, with cluster 6 always being the most distinct, and cluster 0 being central (**Fig. S4B’-D’**).

### 3.2. Marker gene identification provides the first clues for cluster identification

Marker genes were identified for each cluster based on the average fold change value (avgFC) and the adjusted p-value (Padj) (**Table S2**). Results also include the percentage of cells expressing the marker genes in the cluster (pct.1) and in other clusters (pct.2). After filtering (Padj ≤ 0.01 and log_2_(avgFC) ≥ 0.5), we obtained a total of 10,996 marker genes. These were not unequally distributed among the different clusters, with some having only a few marker genes (*e.g.*, cluster 0: only 9 marker genes; 0.81%) and some having a high number of marker genes (*e.g.*, cluster 2: 5,529 marker genes; 49.8%) (**Table S2**). To reveal the best marker genes for each cluster, the five genes with the highest avgFC were selected for visualisation. However, a large proportion was not annotated against the Nr database (56.3%), with some clusters having only not annotated marker genes among their five best markers (*e.g.*, cluster 8), making it difficult to have a first functional interpretation of these clusters. To avoid this, the representation considers only marker genes that were annotated against the Nr database (**Fig. 2A**). The full annotation for these genes, as well as for the unfiltered marker genes list (*i.e.*, considering genes with “NA” as annotation), is provided in **Table S3**.

**Fig. 2.**
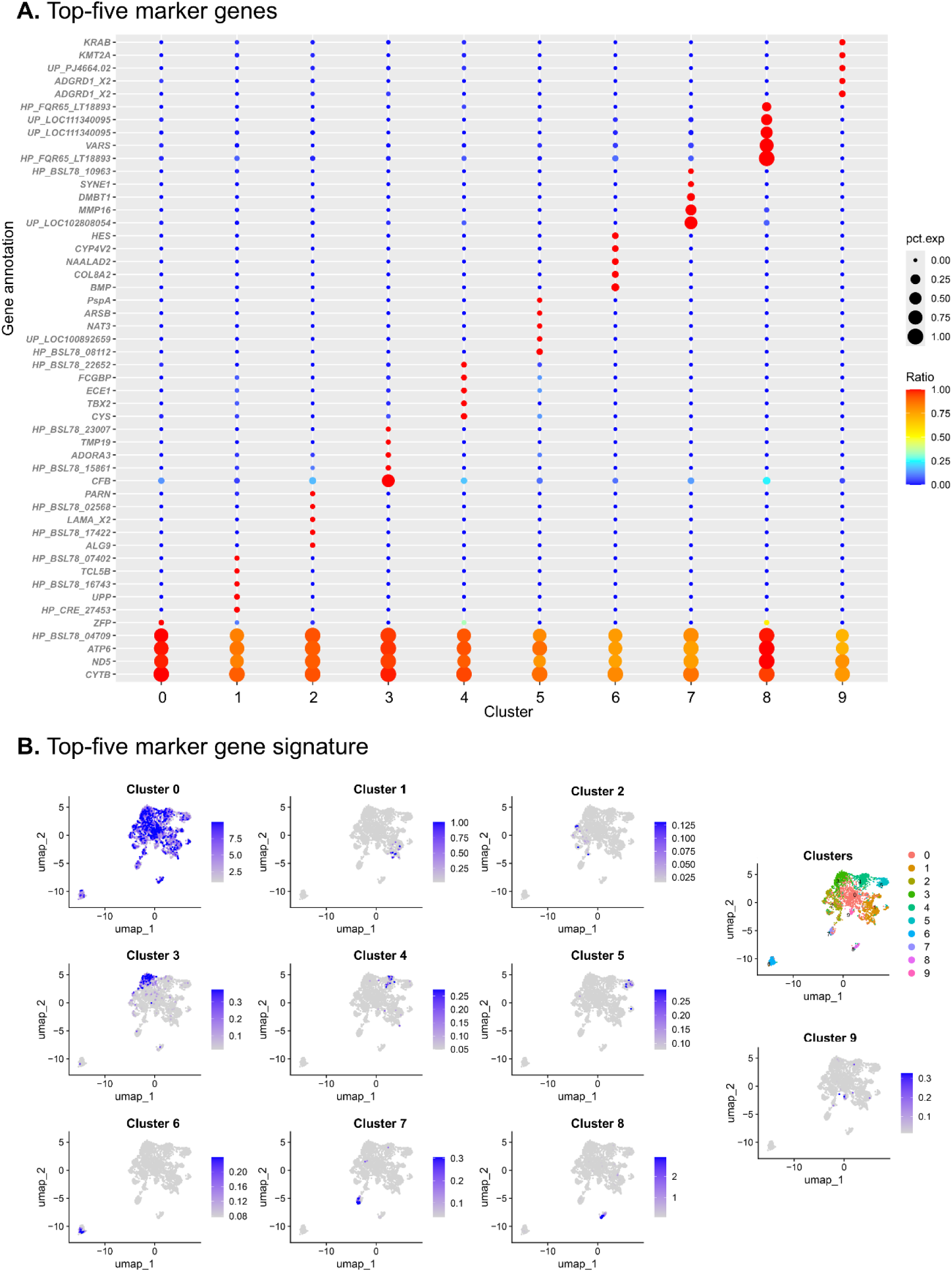
Identification of marker genes for each cluster identified from single-cell RNA sequencing on perivisceral fluid coelomocytes of *Holothuria forskali*. **A.** Five best marker genes for each cluster. Only genes annotated against the Nr database were considered. For each gene, the ratio (colour scale) means the ratio of expression between the cells of the considered cluster and other cells, and the percentage expression (pct.exp) means the percentage of cells expressing the gene within the cluster (full annotation and metrics are shown in **Table S2**). **B.** Expression signature of the five best marker genes (same as in A) visualised in the UMAP. The initial cluster is shown on the upper right. Note that while some signatures are very specific to the cluster (*e.g.,* clusters 6, 7 and 8), some are not specific, especially the signature of cluster 0.

Among genes of interest with regard to the immune function, techylectin-5B is a marker for cluster 1 (*TCL5 B*), complement factor B for cluster 3 (*CFB*), and deleted in malignant brain tumours 1 protein (*DMBT1*) for cluster 7. Lectins are glycoproteins that can link sugar motifs of pathogens and therefore constitute important pathogen receptors (Ren et al., 2025). It was also shown that some have a function of agglutinin in sea cucumbers (Taguchi et al., 2021). *CFB* is an important component of the complement system, participating in the cleavage of C3, essential for the amplification of the signal of opsonisation (Zhong et al., 2012; Xiao et al., 2022). Finally, *DMBT1* is also recognised as an important immune marker gene in echinoderms (Hirano et al., 2016; Ge et al., 2025) due to its scavenger receptor cysteine-rich domain. Expression of these marker genes indicates that clusters 3 and 7 are particularly involved in innate immunity.

Interestingly, a cytochrome P450 4V2 (*CYP4V2*) was identified in the five best marker genes of cluster 6, and three others were in the list of marker genes for this cluster. Cytochrome P450 were reported to be a marker gene for carotenocytes. Although these genes in particular were not those reported to be overexpressed in carotenocytes in Wambreuse et al. (2025a), the presence of marker genes annotated as belonging to the cytochrome P450 family may indicate that this cluster corresponds to cluster 6 (Wambreuse et al., 2025a). In carotenocytes, these proteins are suspected to be involved in carotenoid metabolism, notably by having a bifunctional activity of carotenoid-ketolase, converting beta-carotene into keto-carotenoids (astaxanthin and canthaxanthin) (Mundy et al., 2016; Weaver et al., 2020; Shi et al., 2025). In the same way, a retinol dehydrogenase was identified within the ten best marker genes for this cluster, and two others were identified in the 950 marker genes. These were also identified in bRNA-seq analysis as well as in a proteomics analysis as markers of carotenocytes and are suspected to be involved in carotenoid metabolism, as retinol is a derivative of beta-carotene (Maoka, 2011). Other gene markers for this cluster include the transcription factor “hairy and enhancer of split” (Hes), a bone morphogenetic protein (*BMP*), a collagen alpha-2(VIII) (*COLAA2*), and finally N-acetylated-alpha-linked acidic dipeptidase 2 (NAALAD2).

The remaining marker genes are difficult to interpret as they are either coding for uncharacterised/hypothetical or for non-immune-related proteins. However, it can be highlighted that the marker genes for cluster 0 are also highly expressed in other clusters, and that their annotation corresponds to “housekeeping genes”, including cytochrome B (*CYTB*), NADH dehydrogenase subunit 5 (*ND5*) and ATP synthase F0 subunit 6 (*ATP6*), notably involved in energy metabolism (Da Fonseca et al., 2008). This reflects the low degree of specificity for this cluster, consistent with its position on the UMAP (Fig. 1D). Nonetheless, it is worth highlighting that “energetic metabolism” was reported as a marker of lymphoid-like cells in *A. japonicus* (Yu et al., 2023). This was done by comparing the expression of two coelomocyte subsets by bRNA-seq, among which lymphoid cells correspond morphologically to small round cells in *H. forskali* (Wambreuse et al., 2025a), or “progenitor cells” and “lymphocytes” in other species (Eliseikina and Magarlamov, 2002; Ramírez-Gómez and García-Arrarás, 2010; Caulier et al., 2020). This therefore supports the fact that this cluster corresponds to undifferentiated coelomocytes and is also consistent with the higher proportion of mitochondrial genes expressed in this cluster (**Fig. S2C**)

A signature score was constructed using the “PercentageFeatureSet” function, corresponding to the percentage of total counts in each cluster for a set of genes (corresponding here to the five best marker genes with an annotation). This signature was visualised on the UMAP for each cluster (**Fig. 2B**). Once again, cluster 0 shows a low specificity with a signature observed in all clusters. For other clusters, their signature colocalises with the cluster, consistent with the detection of maker genes. However, some clusters show a lower degree of cells presenting a high signature level, such as clusters 1 and 2. In contrast, clusters 3, 6, 7, and 8 show a higher proportion of cells presenting high signature levels (**Fig. 2B**).

### 3.3. Functional enrichment analyses reveal key immune functions

To gain further functional insights into the different clusters, a functional enrichment analysis was conducted using KEGG and Gene Ontology databases. These analyses yield several metrics, including a fold change enrichment value, which corresponds to the proportion of genes in our dataset for a given functional term relative to the same proportion in a set of random genes. Moreover, a q-value is provided, consisting of an adjusted p-value representing the significance of the enrichment. Given that the number of genes across clusters was highly variable, which may influence the detection of significant enrichment, we present the five best functional entries for each cluster based on fold-change enrichment, regardless of their enrichment significance (**Fig. 3**). However, on **Fig. 3**, a scale colour shows the q-value and the results of the analyses, including all metrics, are shown in **Table S4**.

**Fig. 3.**
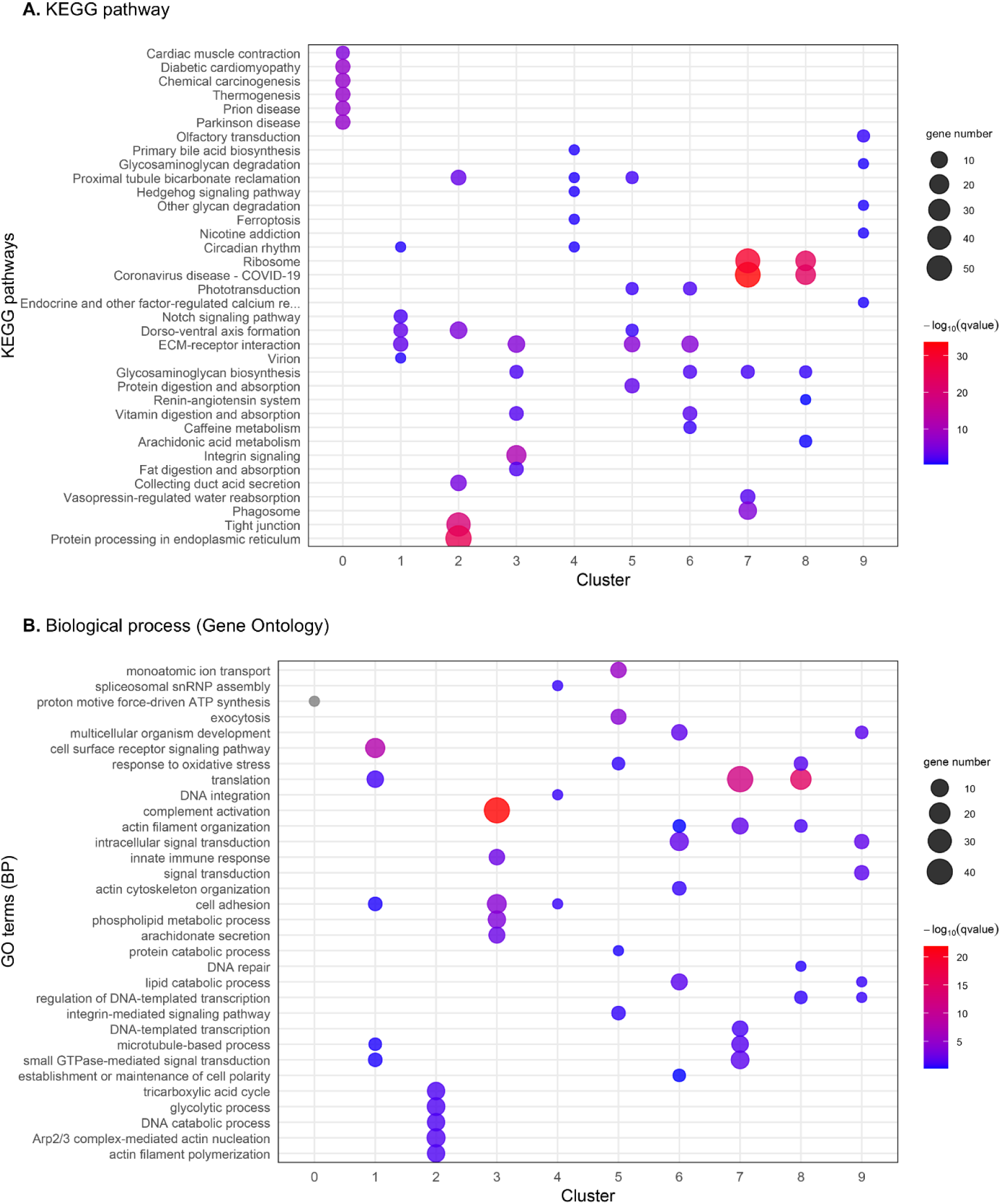
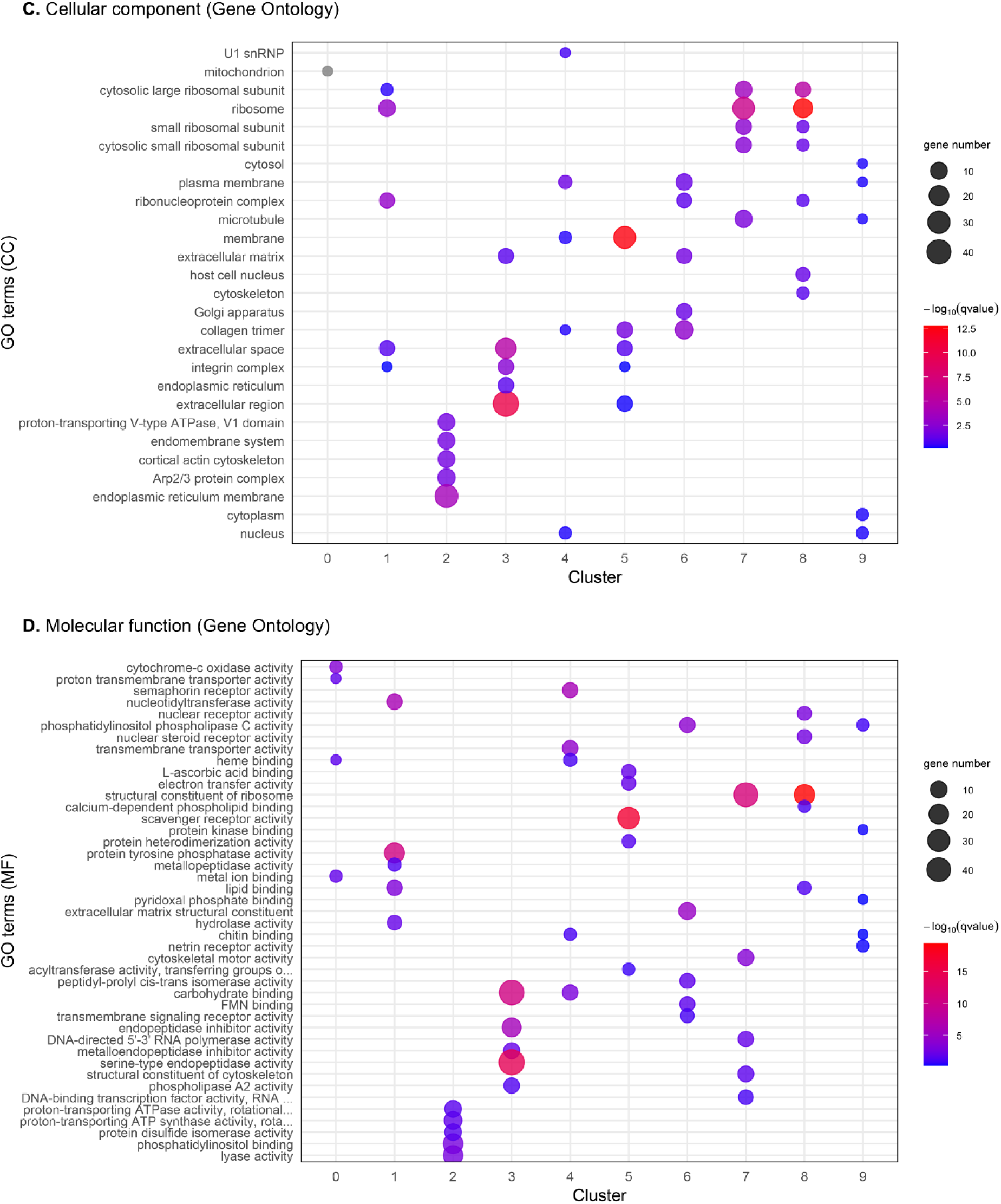
Functional enrichment analysis of cell cluster marker genes identified from single-cell RNA sequencing on perivisceral fluid coelomocytes of *Holothuria forskali*. The five most enriched terms for each cluster are based on the fold enrichment value (note that some clusters have more than five terms for the same analysis; it means that some functional terms were ex-aequo). Dot size corresponds to the number of genes in the pathway, and colour corresponds to the q-value (significance of the enrichment). **A.** Functional enrichment against the KEGG pathway database. **B.** Functional enrichment against the gene ontology database, considering only biological process (BP). **C.** Functional enrichment against the Gene Ontology database, considering only cellular component (CC). **D.** Functional enrichment against the Gene Ontology database, considering only molecular function (MF). Full results of the functional analysis are shown in **Table S4**.

Regarding the KEGG database, a total of 517 terms were significantly enriched, considering all clusters (q-value ≤ 0.05). Four pathways were highly enriched, including “Ribosome”, “Coronavirus disease – COVID-19” in clusters 7 and 8, and “Tight junction” and “Protein processing in endoplasmic reticulum” in cluster 2 (**Fig. 3A**). It can be noted that these pathways include a large number of “generalist” genes, although “Coronavirus disease – COVID-19” contains also many immune genes (see pathway maps: hsa05171). Because these terms are generalist or contain a lot of generalist proteins, enrichment of these pathways should be interpreted with caution. Besides, some interesting pathways related to the immune system could be identified, including “Phagosome” for cluster 7 and “ECM-receptor interaction” for clusters 1, 3, 5 and 6. Looking at gene Nr annotation within the “Phagosome” pathway reveals notably the expression of C-type lectins, complement C3, integrins, V-type proton ATPase and elements of the cytoskeleton.

Functional enrichment against the biological processes of the gene ontology database revealed 21 significant functional enrichments (q-value ≤ 0.05). Among these, the term “Complement activation” of cluster 3 was the most enriched (q = 1.56 × 10^-22^), with 39 genes (**Fig. 3B**). Most of them were either annotated as complement C3 or factor B. Several other biological processes related to the immune system could be identified, including “Innate immune response” for cluster 3, “Response to oxidative stress” for clusters 5 and 8, “Cell adhesion” for clusters 1, 3 and 4 and “Integrin-mediated signalling pathway” for cluster 5. “Innate immune response” comprised genes annotated as “Scavenger receptor cysteine-rich domain protein” (*SRCR*) and “tumour necrosis factor receptor-associated factor 3” (*TRAF3*). In addition, in the pathway “Response to oxidative stress”, six genes were annotated as “dual oxidase 1 precursor” (*DUOX1*) and one as “Catalase” (*CAT*). The most enriched term for cluster 6 was “lipid catabolic process” with mainly genes annotated as “1-phosphatidylinositol 4,5-bisphosphate phosphodiesterase” (*PLCD1*). Cluster 0 has a unique enriched biological process, namely “proton motive force-driven ATP synthesis”.

For the cellular component, 33 terms were significantly enriched. The most enriched terms include “Ribosome” for cluster 8, “Membrane” for cluster 5 and “Extracellular Region” for cluster 3. Among terms of interest, the “Integrin complex is enriched in clusters 1, 3 and 5 (**Fig. 3C**). Regarding cluster 6, four terms were significantly enriched, including “Plasma membrane”, “Extracellular matrix”, “Golgi apparatus” and “Collagen trimer”. Finally, 63 terms were significantly enriched for the molecular function category of the Gene Ontology database. The most enriched terms include “Structural constituent of ribosome” for cluster 8, “Scavenger receptor activity” for cluster 5 and “Serine-type endopeptidase activity” for cluster 3 (**Fig. 3D**).

Enrichment analysis demonstrated that several immune mechanisms were enriched among the different populations of coelomocytes. However, it is still difficult to associate these specific immune mechanisms with specific cell morphotypes due to the lack of functional information in the literature.

The results obtained for cluster 0 are consistent with their low number of marker genes, resulting in no significant enrichment for some datasets, including biological process and cellular component (**Table S3**). Some other terms may suggest that some clusters correspond to phagocytes, which were the most represented cell types in the processed sample. Cluster 7 has, for example, “Phagosome” as a significant enrichment. In addition, the term “Complement activation” was highly enriched in cluster 3, among which several genes encoded the complement C3. This gene is the central element of the complement system, which is thought to have a function in opsonisation, as in vertebrates, and therefore promotes phagocytosis (Smith et al., 2023). In sea urchins, immunocytochemistry and RT-qPCR analyses have revealed that the phagocyte subset was the one having the highest expression of *C3* (Gross et al., 2000). In this way, cluster 3 also constitutes a good candidate cluster for phagocytes.

Clusters 5 and 8 had “Response to oxidative stress” as an enriched term, which may reflect their function in the production of reactive oxygen species (ROS). A recent analysis suggested that phagocytes and spherulocytes are the main ROS-producing cells, which may be consistent with these two clusters (Wambreuse et al., 2025a). Several other immune terms were enriched, such as the term “innate immune response”, which comprises *SRCR* genes for cluster 3. The term “Scavenger receptor activity” for cluster 5 was also enriched for cluster 5 in the molecular function database. *SRCR* are important pathogen receptors in invertebrates (Peng et al., 2024), highlighting the function of this cluster as immune sentinels. *TARF3* was also a marker of cluster 3 among the “Innate immune response pathway”. This gene is an adaptor protein that regulates innate immune signalling, particularly NF-κB–mediated inflammation (Yang et al., 2016), demonstrating, again, the key function of cluster 3 in immunity.

Based on these enrichments, several clusters may be associated with phagocytes. However, this is not unexpected for several reasons. Firstly, none of the clusters matches the proportion of phagocytes observed in the processed sample. Secondly, multiple phagocyte morphotypes have been described in sea cucumbers, including petaloid and filiform phagocytes (Hetzel, 1963; Chia and Xing, 1996; Wambreuse et al., 2025a), which probably have distinct gene expression. For example, in sea urchins, different phagocyte types are distinguished based on their shape and the structure of their cytoskeleton, and these show distinct gene expression patterns, including, for example, within the transformer gene family (Smith and Lun, 2017). It should therefore not be surprising to have several clusters corresponding to phagocytes, and clusters 3, 5 and 7 are the best candidates based on the functional enrichment.

Regarding enrichment for cluster 6, which is suspected to be carotenocytes, the most enriched KEGG pathway was “ECM-receptor interaction” (q-value = 1.15 × 10^-6^). This could be consistent with the ability of these cells to remain marginated (Caulier et al., 2020; Wambreuse et al., 2025a), *i.e.*, attached to the wall of tissues rather than freely floating in the lumen. This is also supported by the molecular function database, for which the term “extracellular matrix structural constituent” was highly enriched (q-value = 1.33 × 10^-5^). Regarding the biological process database, three terms were significantly enriched, including “intracellular signal transduction”, “multicellular organism development” and “lipid catabolic process”. This last was also reported as an enriched term in carotenocytes in Wambreuse et al. (2025a) and may reflect their function in carotenoid storage, as carotenoids are lipidic molecules. However, most genes enriched for this pathway encoded *PLCD1*, whose function is unknown in echinoderms, and the link with carotenoid metabolism is unclear.

Although enrichment analyses are useful to provide a broad function of the cluster, they should not be taken literally, as they are mainly based on vertebrate and especially human physiology. Looking at the genes behind the terms is therefore crucial to validate some putative functions, and further analysis should aim to validate the expression of some interesting genes within the different morphotypes. Previous enrichment analyses have been performed on putative coelomocyte clusters from the regenerating intestine of *H. glaberrima* (Medina-Feliciano et al., 2025). Although the enriched terms differ from those reported here, this discrepancy is likely due to differences in biological context, as those cells were analysed within regenerating tissues, which are expected to exhibit distinct transcriptional programs.

### 3.4. Differential gene expression from bRNA-seq allows the identification of the carotenocyte cluster

Cell counting of the sample processed in scRNA-seq showed the presence of carotenocytes – small red cells rich in carotenoids (**Fig. 1B** and **C**). Previous work carried out by our laboratory showed that these cells are highly present in the hydrovascular fluid (HF), in contrast to the perivisceral fluid (PF). By comparing the expression profile between the two fluids, it was possible to obtain a list of candidate marker genes for carotenocytes in particular (2,070 genes) versus all other cell types (783 genes) (see Wambreuse et al., 2025a). In this section, these two gene lists are referred to as the carotenocyte-enriched sample (CAE – 80.6 ± 5.1% of carotenocytes; n = 6) and the other coelomocyte-enriched sample (OCE – 0.75 ± 0.71% of carotenocytes; n = 6; **Fig. 4A**). In our clustering analysis, a cluster was highly divergent from the others, corresponding to cluster 6 and appeared to express some genes characteristic of carotenocytes. As visual examination of the sample revealed 7.0% of carotenocytes in the perivisceral fluid sample, it was postulated that this small cluster, composed of 4.7% of the cells, may correspond to carotenocytes.

**Fig. 4.**
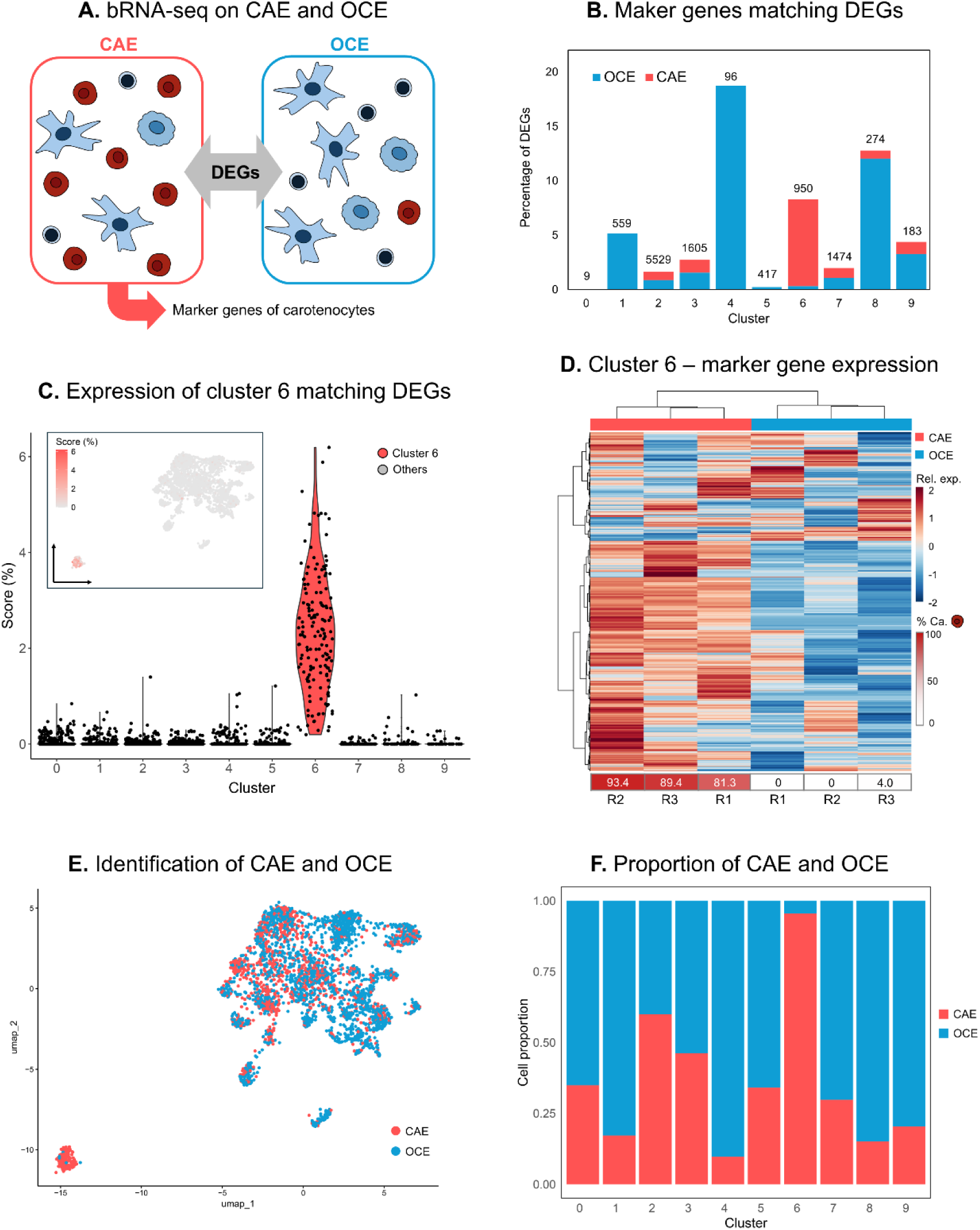
Identification of carotenocytes based on differential expression between enriched coelomocyte populations by bulk RNA sequencing (bRNA-seq) in *Holothuria forskali*. **A.** Illustration of differential expression analyses perivisceral fluid (containing almost no carotenocytes; called other coelomocyte enriched samples – OCE) and hydrovascular fluid (highly enriched in carotenocytes; called carotenocyte-enriched samples – CAE). Among differentially expressed genes (DEGs), up-regulated genes in CAE are presumed to be markers for carotenocytes. **B.** Proportion of marker genes in each cluster matching DEGs, either up-regulated in OCE or in CAE (in blue and red, respectively; total number of marker genes per cluster is indicated). **C.** Expression signature of the overexpressed genes in CAE matching cluster 6 marker genes. This expression is mapped on the UMAP and compared between clusters using a violin plot. **D.** A heatmap representing the expression (autoscalled log(FPKM)) of cluster 6 marker genes (950 genes) between CAE and OCE samples. The level of enrichment of carotenocytes in each sample is shown as a percentage below the heatmap. **E.** Identification of CAE and OCE positive cells using SingleR based on the mean expression level of DEGs between CAE and OCE (2,853 genes). **F.** Proportion of cells identified as OCE or CAE cells. Cluster 6 shows more than 95% of CAE cells.

To further confirm this correspondence with a larger gene panel, a first analysis was conducted to count the number of marker genes that match differentially expressed genes (DEGs) between CAE and OCE, and among these, which ones were overexpressed in CAE or OCE. This number was converted as a percentage of marker genes in each cluster (**Fig. 4B**). While cluster 4 has the highest proportion of marker genes (18,7%), all of them were overexpressed in OCE. Moreover, all clusters had a proportion of marker genes overexpressed in OCE higher than in CAE, except cluster 6, which had 80 marker genes matching the differential expression (8.32% of their marker genes), including 76 that were overexpressed in CAE. To further analyse the expression of these genes, a gene signature score was built using the “PercentageFeatureSet” function, based on the 76 marker genes, and the signature score expression was compared between clusters and visualised on the UMAP (**Fig. 4C**). This signature is clearly localised on cluster 6, with good homogeneity within the cluster. Moreover, with regard to the violin plot, cluster 6 cells had an overall higher score, supporting the hypothesis that this cluster corresponds to carotenocytes (**Fig. 4C**).

To test the hypothesis using another approach, the expression of the 950 marker genes of cluster 6 was compared between CAE and OCE samples using a heatmap (**Fig. 4D**). It revealed a good clustering between the two types of samples, with overall higher expression of the marker genes in CAE samples. Carotenocytes proportion in each sample was added beyond the heatmap to show the level of enrichment in this cell population in CAE and OCE (88.0 ± 6.1% in CAE against 1.3 ± 2.9% in OCE; n = 3).

Finally, we used the tools SingleR to attribute each cell of our dataset either to the identity of CAE or OCE based on the mean expression of the 2853 DEGs within the two groups. While CAE-positive cells could be observed within other clusters, cluster 6 appears to be almost only composed of CAE-positive cells (**Fig. 4E**). The proportion of the two types of samples was visualised for each cluster and cluster 6 was highly positive to CAE with 95.8% of cells identified as such (**Fig. 4F**). Most of other clusters had a proportion of CAE below 50%, except cluster 2 for which this proportion reached 59.1%. A possible explanation for this may be that cluster 2 was pointed out as the one having the highest proportion of multiplets. Therefore, it could be hypothesised that some cells are doublets with carotenocytes.

### 3.5. ScRNA-seq provides novel insight into carotenocyte expression

Carotenocytes have been recently discovered as a new type of pigmented coelomocytes in echinoderm (**Fig. 5A**). Different molecular data could be acquired using comparative transcriptomics and proteomics for these cell types (Wambreuse et al., 2025a). However, these data were acquired on an enriched population of carotenocytes and not directly on a pure population, constituting an approximation of the real gene expression of these cells. Comparing the scRNA-seq datasets with the list of candidate marker genes previously obtained allows us to find 76 genes in common between the two analyses, which can be considered as highly confident marker genes for carotenocytes (**Fig. 5B**). While this number can seem low, it is important to note that scRNA-seq does not have the same sensitivity as bRNA-seq, meaning that genes having low expression could not have been detected. In addition, although the 950 marker genes of cluster 6 were not all statistically differentially expressed between OCE and CAE, they showed an overall higher expression in CAE, supporting the fact that these genes could also be considered as carotenocyte-specific marker genes.

**Fig. 5.**
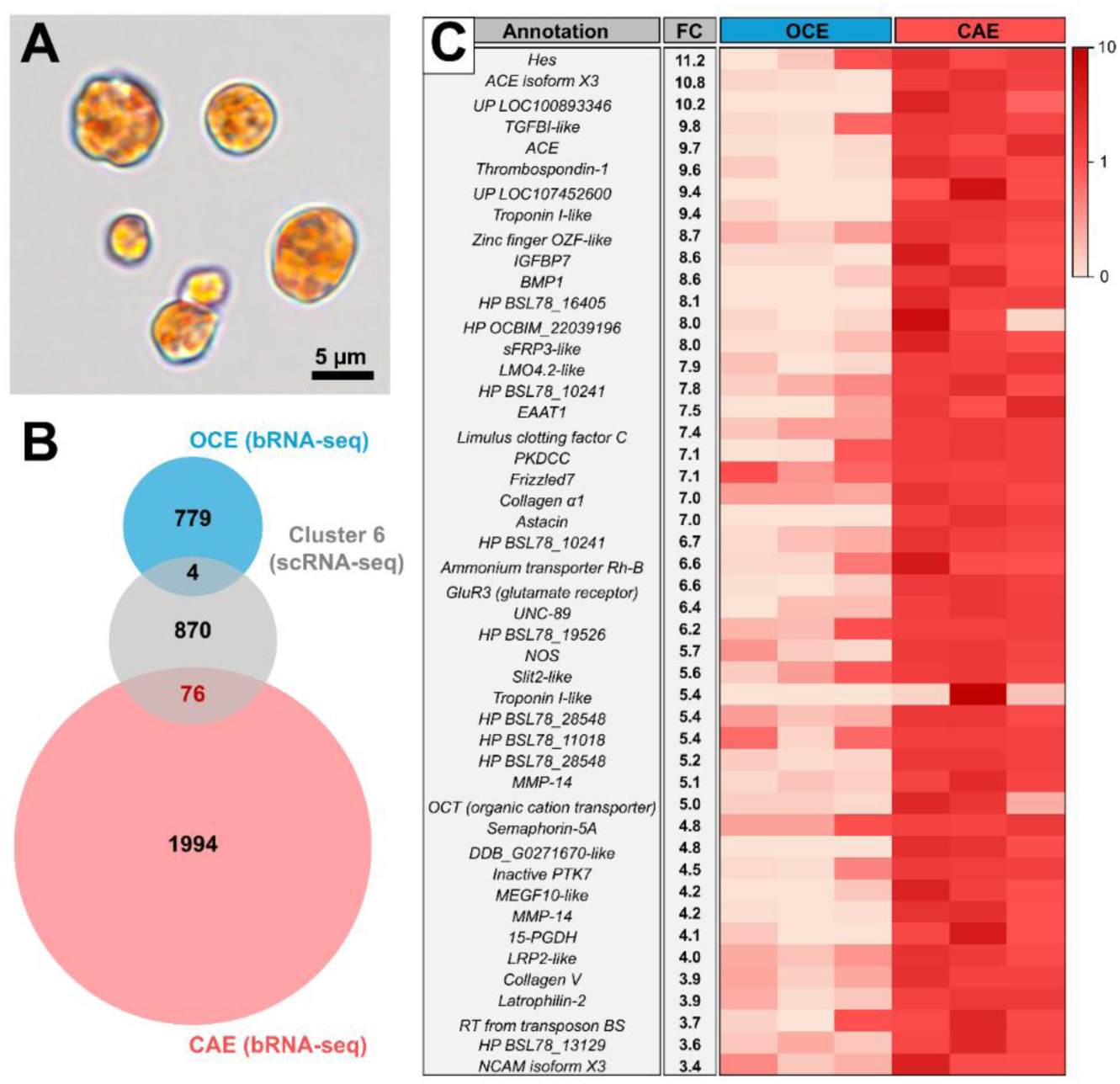
List of marker genes identified for carotenocytes in *Holothuria forskali* based on consistency between bulk and single-cell RNA sequencing analyses. **A.** Picture of several carotenocytes from the hydrovascular fluid of *H. forskali*. **B.** Venn diagram illustrating the selection of marker genes made by comparing the single-cell RNA sequencing (scRNA-seq) and bulk RNA sequencing (bRNA-seq) datasets. The selected genes are the common ones between genes overexpressed in the caerotenocytes-enriched samples (CAE – bRNA-seq), marker genes of cluster 6 (Cl.6 – scRNA-seq) identified as the carotenocyte cluster. It can be seen that the other coelomocyte-enriched samples (OCE – bRNA-seq) and cluster 6 share only a few genes. **C.** List of selected genes with their annotation, scRNA-seq average fold change value (average log_2_(fold-change) – FC) and expression (autoscalled log(FPKM+1)) in three samples of CAE and OCE (only the annotated genes among the 76 genes are represented; the full table, including the complete annotation, can be found in **Table S5**).

To obtain more functional information on carotenocytes, it can be useful to look at the annotation of these 76 highly confident marker genes. Among these, only 47 could be annotated against the Nr database. The expression of these genes in OCE and CAE samples is represented in **Figure 5C**, along with their annotation. These were ordered according to their scRNA-seq avgFC, reflecting the specificity for the carotenocyte cluster. The best marker gene was a transcription factor, namely “hairy and enhancer of split” (*Hes*). Interestingly, this gene was the only one annotated as *Hes* in the transcriptome of *H. forskali*. *Hes* transcription factor is known to be a downstream gene of Notch signalling activation, which regulates cell differentiation through cell-cell interaction (Iso et al., 2003). More specifically, *He*s acts as a repressor of other transcription factors that induce cell differentiation and is therefore considered to have a function in stem cell maintenance (Kobayashi and Kageyama, 2014). In fruit flies and humans, this transcription factor is known to play a crucial role in neuronal development (Kobayashi et al., 2014). Interestingly, in sea urchin embryos, perturbations of its expression resulted in a higher number of pigmented cells (Erkenbrack, 2018), suggesting that this transcription factor may be involved in pigment cell differentiation in sea cucumbers as well, although the embryonic origin of carotenocytes is still totally unknown.

Among marker genes, several proteins involved in cell adhesion and the production of extracellular matrix can be identified, including limulus clotting factor C, transforming growth factor-beta-induced protein ig-h3-like (*TGBBI-like*), LIM domain transcription factor LMO4.2-like (*LMO4.2-like*), and collagens. Limulus clotting factor C was initially identified in horseshoe crabs, in which it is produced by their immune cells and has a function in binding lipopolysaccharides, *i.e.,* bacterial endotoxin, and initiates the coagulation cascade (Wang et al., 2003). *TGBBI-like* and *LMO4.2-like* seem to share sequence similarities, as one gene shares the two annotations. In vertebrates, *TGBBI-like* is an extracellular protein associated with functions including cell adhesion, migration, proliferation and differentiation in diverse cell types (Kim et al., 2009). Interestingly, it was demonstrated that this protein is present in vertebrate platelets and can be released upon activation. These have the effect of activating other platelets and promoting the production of collagen fibres in surrounding tissues, leading to thrombus formation (Kim et al., 2009). This is consistent with the expression of collagen observed in carotenocytes and their ability to remain marginated at the surface of hydrovascular tissues (Wambreuse et al., 2025a). Maker gene list also includes several genes involved in neural cell differentiation and development, such as neuronal cell adhesion molecule (*NCAM*), semaphorin and SLIT homolog 2 protein (*SLIT2*) (Walsh and Doherty, 1996; Itoh et al., 1998; Yazdani and Terman, 2006), which could reflect a function in neural regeneration.

One important category of marker gene for carotenocytes was also Bone Morphogenetic Proteins (*BMPs*), with one gene that was in the top five of marker genes and another that was found in the list of 76 common marker genes. Interestingly, in sea urchin embryos, *BMP2/4* is expressed in primary pigment cells (Duboc et al., 2010). In addition, silencing *BMP2/4* resulted in albino embryos, demonstrating the importance of these genes in the development of pigmented cells in echinoids. While caution should be taken when comparing pigmented coelomocytes between echinoids and holothuroids, which contain different pigment types, the common expression of *BMPs* highlights a putative homology in the function of these proteins in pigmentation of Echinozoa, as well as a putative common origin between carotenocytes and sea urchin pigment cells. However, it should be noted that Perillo et al. (2020) reported several families of genes involved in the production of naphthoquinones, *i.e.,* sea urchin pigment, specific to pigmented circulating cells in sea urchins, that could not be detected in the marker genes of carotenocytes. While genes specific to the metabolism of the distinct pigment may precede the divergence between holothuroids and echinoids, further analyses, notably integrating multi-species datasets, may help to highlight some potential cell homologies.

Finally, a nitric oxide synthase (NOS) was identified among the 76 marker genes. Previous work suggested that carotenocytes represent an antioxidant-rich population, expressing catalase and containing potent antioxidant carotenoids (Wambreuse et al., 2025a). Although catalase was not identified among the marker genes of cluster 6, it can be hypothesised that carotenocytes preferentially rely on nitric oxide (NO) production via NOS as an antimicrobial mechanism (Shao et al., 2016). NO and derived reactive nitrogen species can contribute to pathogen control while potentially limiting reliance on ROS-based mechanisms, which may be rapidly quenched in antioxidant-rich cellular environments (Tan et al., 2020).

Annotations of remaining marker genes were difficult to interpret without molecular studies. However, this list represents a good set of genes to pave the path for further analysis, notably in trying to target carotenocytes with specific molecular markers. Overall, it could also be observed that carotenocytes were transcriptionally highly distinct from the rest of the coelomocytes, which raises interesting questions on the origin of these cells.

### 3.6. First step toward a molecular characterisation of coelomocytes

Using scRNA-seq on a pooled population of coelomocytes encompassing multiple cell morphotypes allowed us to uncover key transcriptomic specificities, highlighting the molecular distinctiveness of immune cell types in holothuroids and the importance of considering these cells as discrete entities in molecular studies. This work represents a first pilot application of single-cell transcriptomic approaches to circulating coelomocytes in holothuroids, a group for which transcriptomic resources on immune cells remain extremely limited. As such, further work will be required to address several limitations inherent to this initial proof-of-concept study.

On one hand, the experiment was conducted on a single biological sample, which may result in a strong individual effect and limit the assessment of inter-individual variability. In addition, data processing was deliberately conservative, with minimal filtering applied. This choice was motivated both by the relatively low number of recovered cells and by the exploratory nature of the study, with the aim of retaining as much biological information as possible. Consequently, no filtering was applied to remove potential doublets or cells expressing a high proportion of mitochondrial genes, parameters known to influence scRNA-seq datasets (McGinnis et al., 2019; Osorio and Cai, 2021).

On the other hand, scRNA-seq reads were mapped to a *de novo* transcriptome, which, although highly complete based on BUSCO scores (Wambreuse et al., 2025a), lacks the structural resolution of a reference genome, generally preferred for single-cell analyses. This approach likely inflated the number of detected genes due to transcript fragmentation and redundancy within the predicted gene catalogue, potentially leading to overrepresentation or duplication among marker genes. Nevertheless, the use of a *de novo* transcriptome ensured consistency with previous differential expression analyses and enabled meaningful functional insights in the absence of a reference genome for this non-model organism.

Taken together, this pioneering study provides a valuable foundation for future investigations of coelomocyte diversity. Further work will be necessary to validate the correspondence between transcriptional clusters and morphological cell types. This could be achieved through bRNA-seq of enriched cell populations, as performed for carotenocytes, or through spatial validation of marker gene expression using antibodies or *in situ* hybridisation. In addition, trajectory analyses may help determine whether coelomocyte morphotypes such as phagocytes, spherule cells, and carotenocytes share a common circulating precursor and clarify the identity and role of putative progenitor cells, potentially corresponding to cluster 0 in this study. Overall, despite its exploratory nature, this first scRNA-seq analysis provides rare and valuable insights into the transcriptomic diversity of echinoderm coelomocytes and offers novel functional information on carotenocytes, a cell type that has only recently been identified.

## 4. Conclusion

This pioneering study employed, for the first time, scRNA-seq on circulating coelomocytes of a holothuroid, enabling the identification of ten distinct transcriptional populations. Among these, several clusters display key immune-related genes, reflecting functional specialisation among coelomocyte types. The central position of cluster 0 within the transcriptional landscape, together with its non-specific gene expression profile, suggests that it may correspond to a “progenitor” cell population. In addition, the use of a gene list from a previous bRNA-seq study on carotenocytes allowed their identification among specific clusters, providing new insights into their gene expression profiles. Although this initial dataset will require validation across additional coelomocyte types, it represents the first characterisation of gene expression in circulating coelomocytes of holothuroids at the single-cell level. Overall, this study constitutes an important first step toward the establishment of a molecular classification of coelomocytes and will be of great interest for comparative immunology within the deuterostome lineage.

## 5. Acknowledgements

The authors would like to thank the collection service of the Roscoff Biological Station for supplying the biological material necessary for this study. They also acknowledge the Genomic Platform of the GIGA Institute for their help with the acquisition of single-cell sequencing data. They are also grateful to Prof. Katherine Buckey, Prof. Patrick Flammang, Prof. David Gillan, Dr Fanny Gaillard and Estelle Bossiroy for their reading and constructive comments on this study. Finally, they would like to extend their thanks to Nathan Puozzo and Antoine Batigny for their logistical help.

## 7. Data availability

Raw data will be made available upon publication of the manuscript. However, some data may be shared by the authors upon reasonable request. The Seurat object can be shared within a collaborative framework.

## 8. Author contribution

Conceptualisation: N.W., L.F., F.B., L.Z., B.D., G.C., I.E., J.D.

Methodology: N.W., A.L., L.F., J.D.

Formal analysis: N.W., J.D.

Investigation: N.W., J.D.

Software: A.L.

Data curation: N.W., A.L.

Writing – original draft: N.W.

Writing – review & editing: A.L., L.F., L.Z., B.D., G.C., I.E., J.D.

Visualisation: N.W.

Funding acquisition: F.B., G.C., I.E., J.D.

Project administration: F.B., I.E.

Supervision: G.C., I.E., J.D.

## 9. List of supplementary

### Supplementary figures

**Fig. S1.** Quality controls and principal component analysis (PCA) on the single-cell RNA sequencing data.

**Fig. S2.** Doublet detection using DoubletFinder.

**Fig. S3.** Detection of mitochondrial gene expression.

**Fig. S4.** Effect of different filtering parameters on the UMAP configuration.

### Supplementary tables

**Table S1.** Gene expression across the dataset.

**Table S2.** List of marker genes of each cluster.

**Table S3.** List of the five marker genes for each cluster.

**Table S4.** Functional enrichment analysis results.

**Table S5.** List the 76 carotenocyte marker genes common between bulk RNA sequencing and single-cell RNA sequencing.

## 9. Supplementary figures

**Fig. S1.**
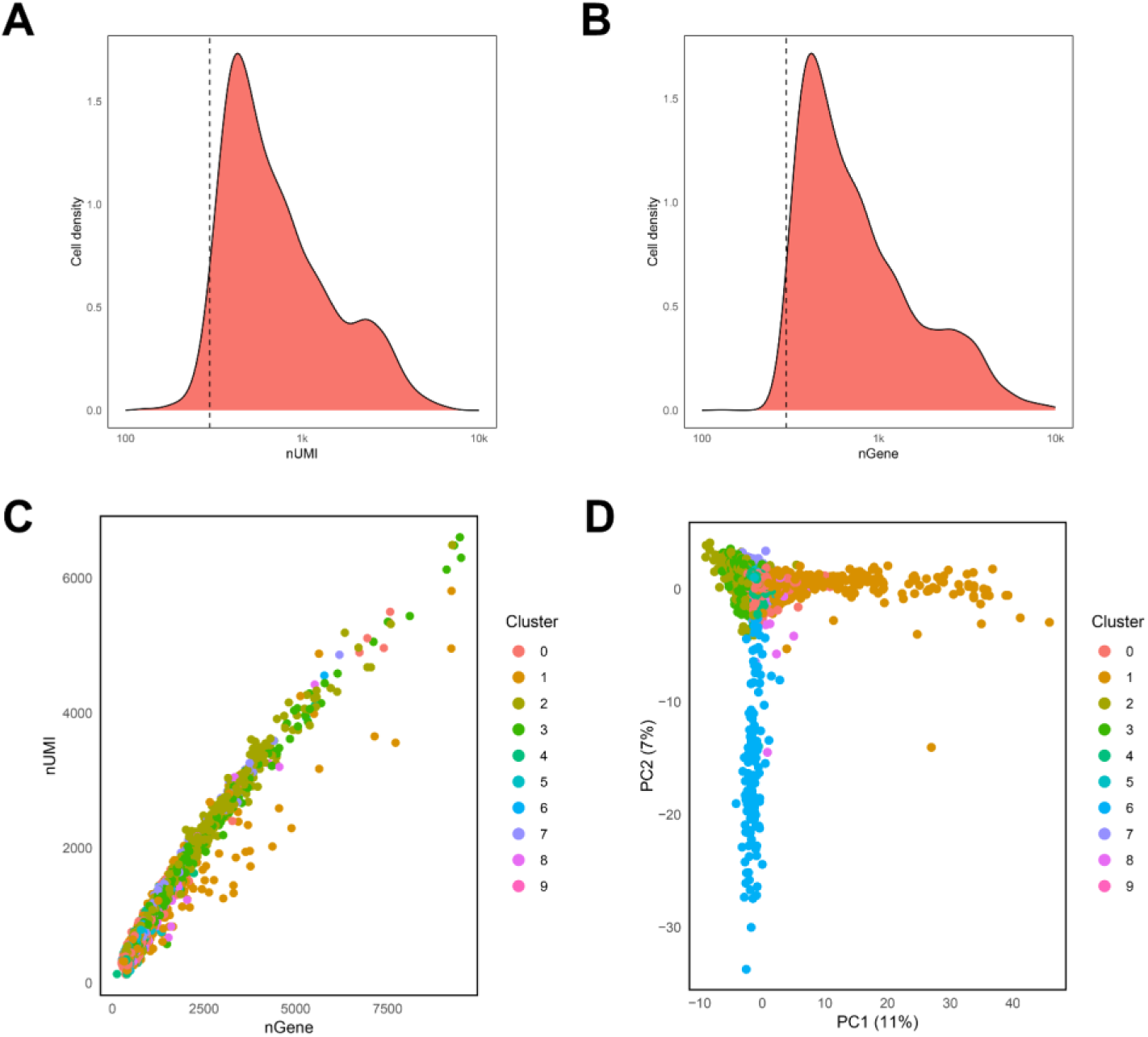
Quality control and principal component analysis (PCA) on the single-cell RNA sequencing data obtained from single-cell RNA sequencing analysis on perivisceral fluid coelomocytes of *Holothuria forskali*. **A.** and **B.** The cell distribution of the number of RNA (UMIs) and the number of genes (Features), respectively. The dotted line corresponds to the 300 UMIs and Features per cell; note that the majority is above this number, reflecting the good quality of the data. **C.** Number of UMIs as a function of number of features, showing a good relationship between these two variables. **D.** PCA on normalised single-cell sequencing data; note that PC1discriminates cluster 2 while PC2 discriminates cluster 6.

**Fig. S2.**
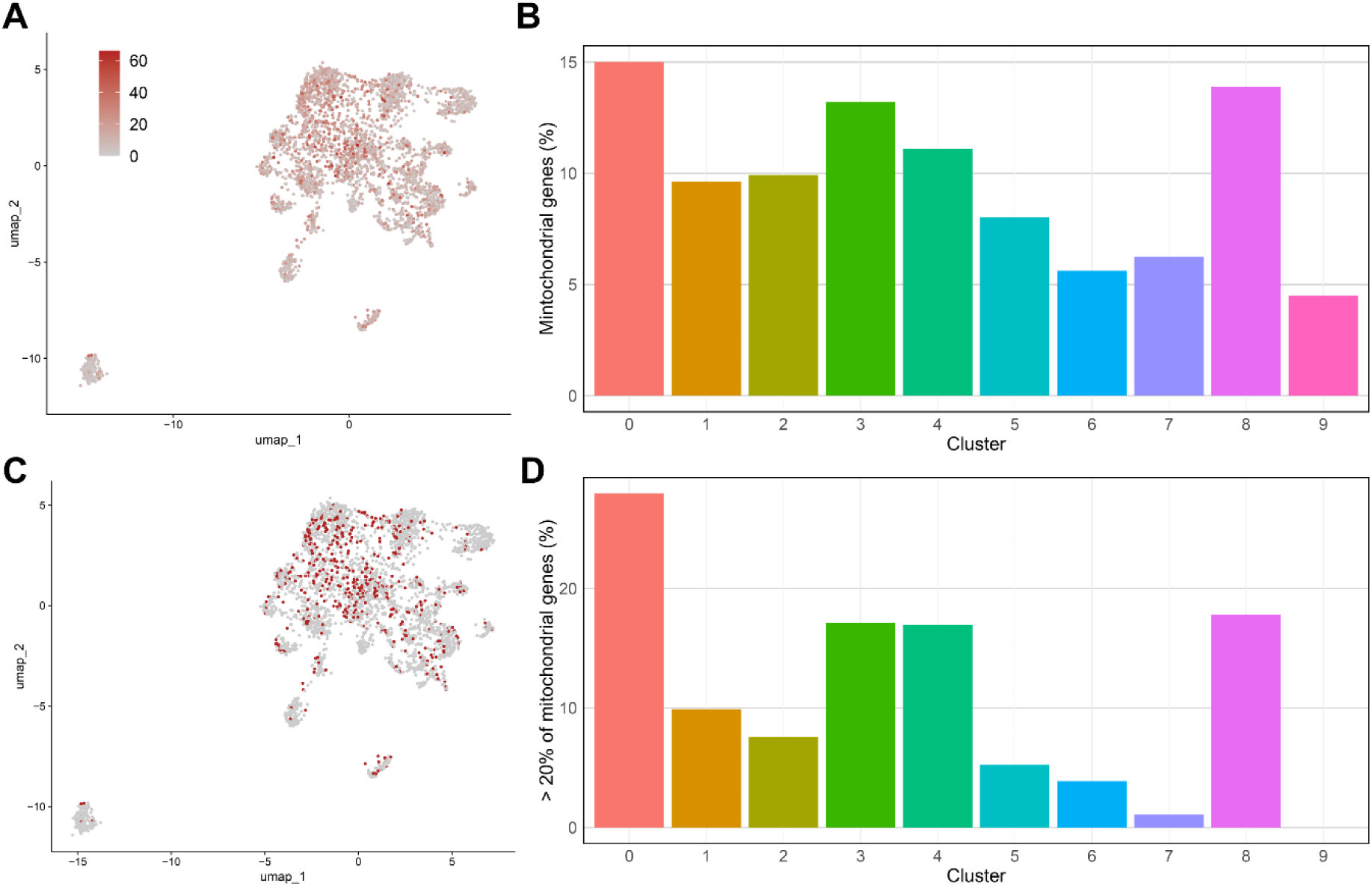
Detection of mitochondrial gene expression on the dataset obtained from single-cell RNA sequencing analysis on perivisceral fluid coelomocytes of *Holothuria forskali*. **A.** Mapping of the percentage of mitochondrial genes across the UMAP. **B.** Mean mitochondrial gene expression in each cluster. **C.** Mapping of cells having a percentage of mitochondrial gene expression > 20%. **D.** Proportion of cells having a percentage of mitochondrial gene expression > 20% in each cluster.

**Fig. S3.**
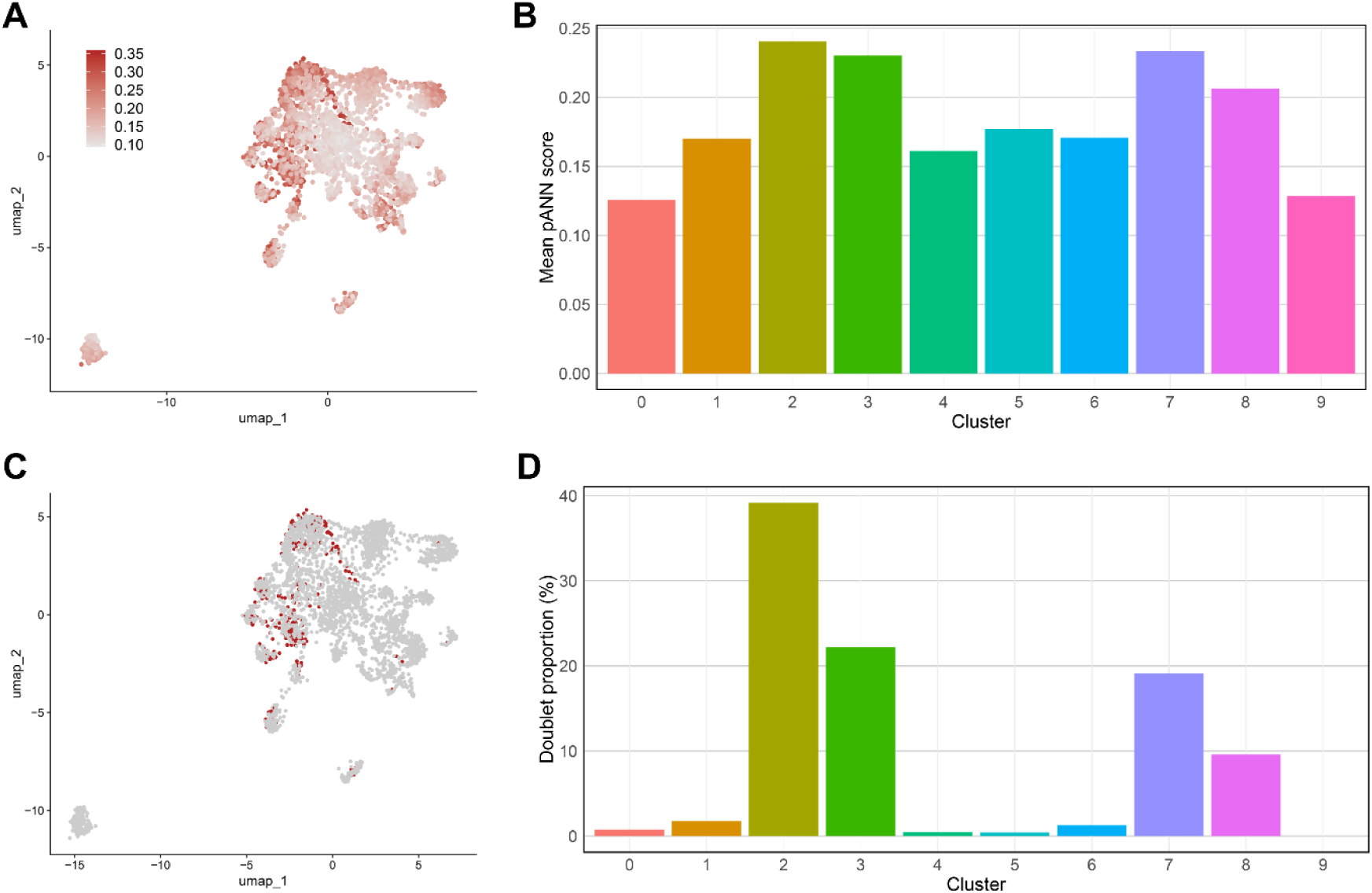
Doublet detection using DoubletFinder on the dataset obtained from single-cell RNA sequencing analysis on perivisceral fluid coelomocytes of *Holothuria forskali*. **A.** Mapping of the paNN score across the UMAP; a higher score means a higher chance of being a doublet. **B.** Mean paNN score for each cluster. **C.** Mapping of cells considered as doublets, *i.e.*, 10% of the cells with the highest paNN score. **D.** Proportion of doublets in each cluster.

**Fig. S4.**
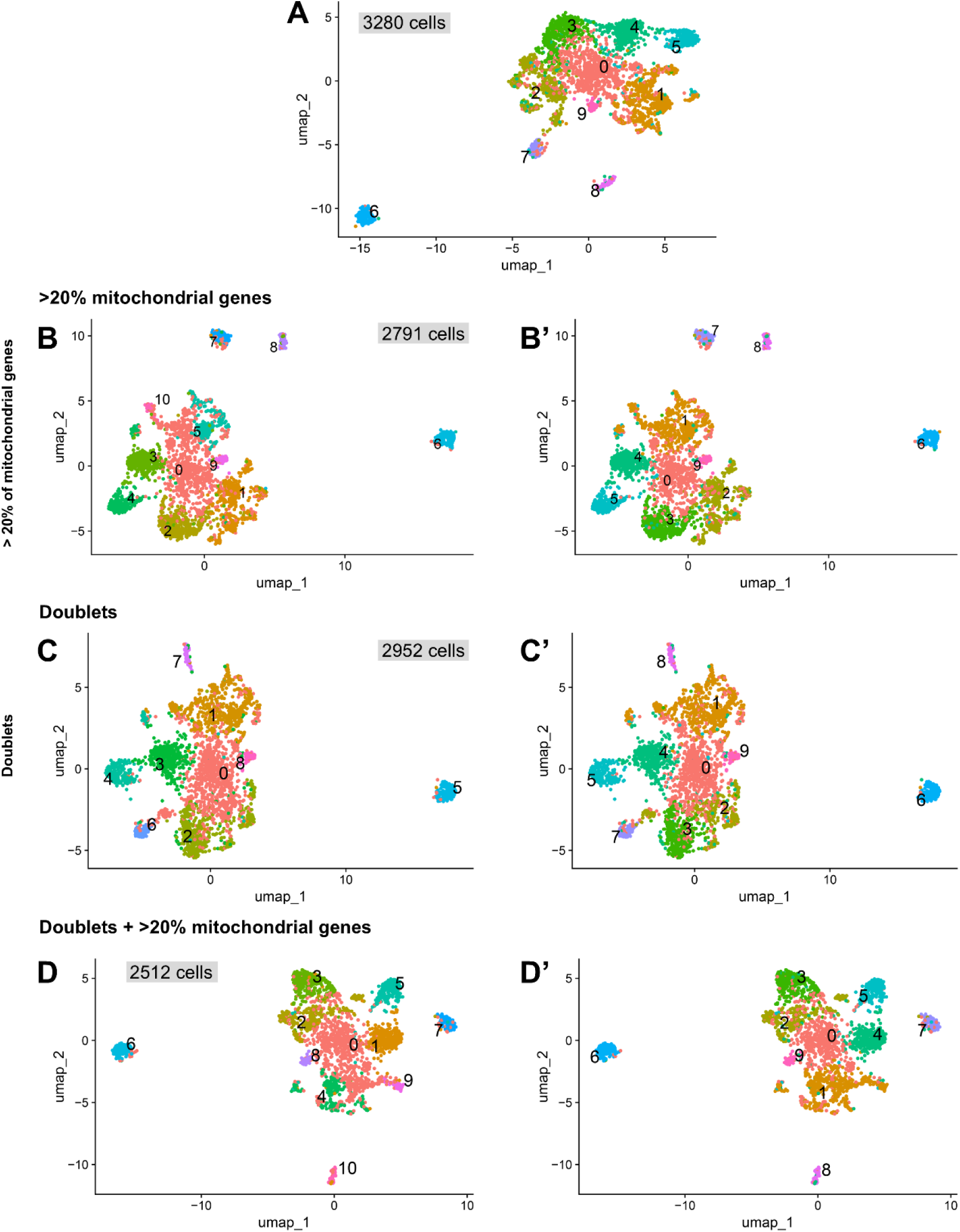
Effect of different filtering parameters on the configuration of the UMAP obtained from single-cell RNA sequencing analysis on perivisceral fluid coelomocytes of *Holothuria forskali*. **A.** Original UMAP with no filtering. **B.** Filter applied on cells that express more than 20% of mitochondrial genes. **C.** Filter applied cells that were identified as doublets using DoubletFinder – 10% cells with the highest paNN score. **D.** Filter from B and C applied together. B’-D’. Corresponding to the UMAP, respectively, but with a mapping of the cluster from the original UMAP (in A). Overall, it can be seen that while some clusters are merged or divided, the general configuration stays similar, including cluster 0 as a central cluster and cluster 6 as the most divergent.

## 10. Supplementary tables

**Table S1.** Average expression (avg) and percentage (pct) of expression for all genes of the dataset in each cluster. The average expression is calculated for all cells included in the considered cluster. The percentage of expression is the number of cells expressing the gene out of the total number of cells in the considered cluster. See the dedicated Excel file.

**Table S2.** List of marker genes of each cluster. Only marker genes with p.adj ≤ 0.01 and log_2_(avgFC) ≥ 0.5 were considered. Annotations against several databases are also provided for each marker gene. See the dedicated Excel file.

**Table S3.** List of the five marker genes for each cluster. These include a filtered list according to the lack of annotation and an unfiltered list. Annotations against several databases are also provided for each marker gene. See the dedicated Excel file.

**Table S4.** Functional enrichment analysis table against KEGG pathways, Biological Processes (BP), Cellular Component (CC) and Molecular Function (MF). Terms significantly enriched are in green (q-value ≤ 0.05). See the dedicated Excel file.

**Table S5.** List of marker genes of carotenocytes supported by both bulk RNA sequencing and single-cell RNA sequencing analyses. Annotations are provided against the Nr database as well as the expression level in carotenocyte-enriched samples (CAE) and other coelomocyte-enriched samples (OCE). See the dedicated Excel file.

