## Supplementary figures and images for "Single-cell transcriptomics reveals transcriptional diversity of sea cucumber perivisceral fluid coelomocytes"

### Fig. S1

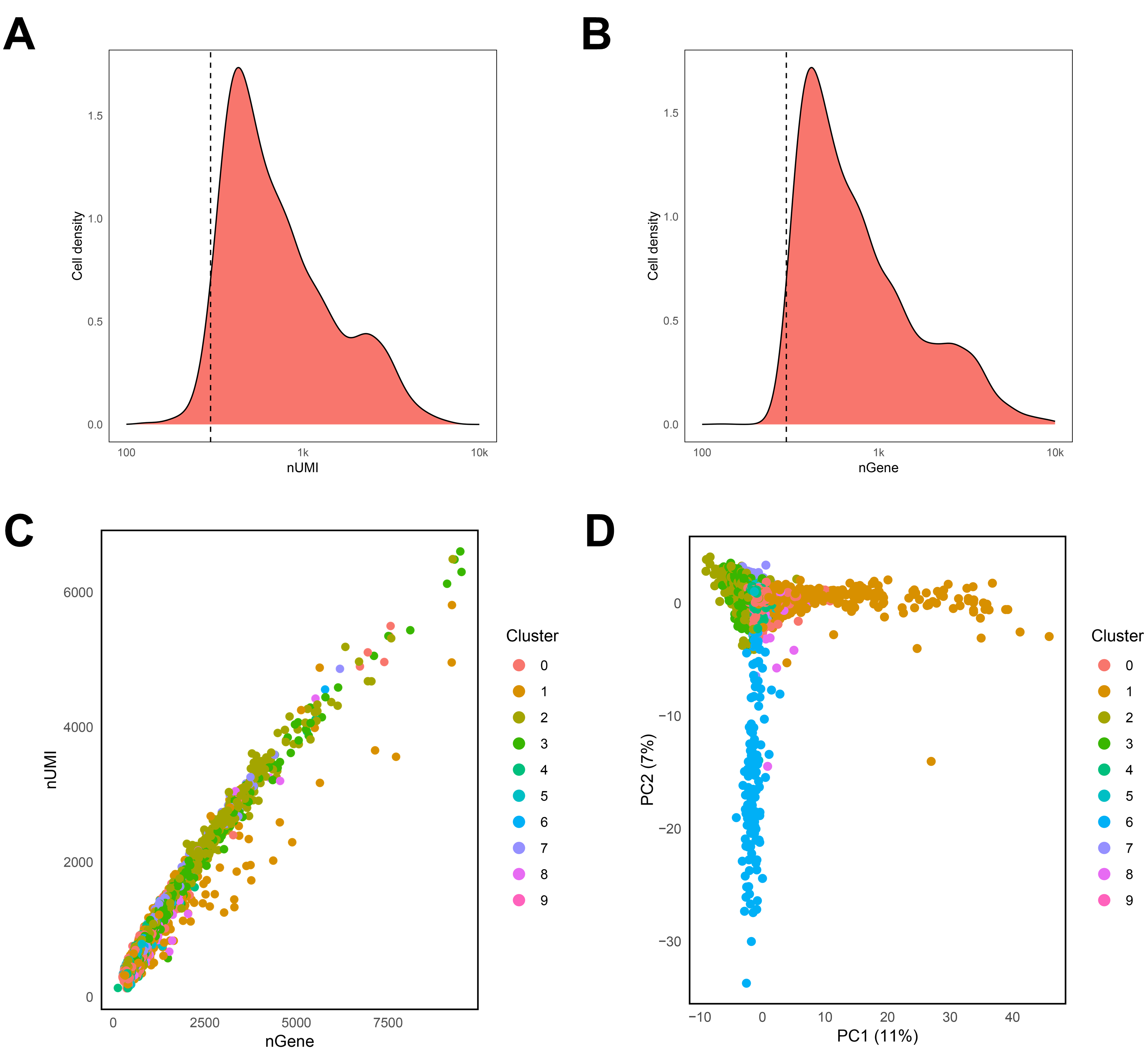

### Fig. S2

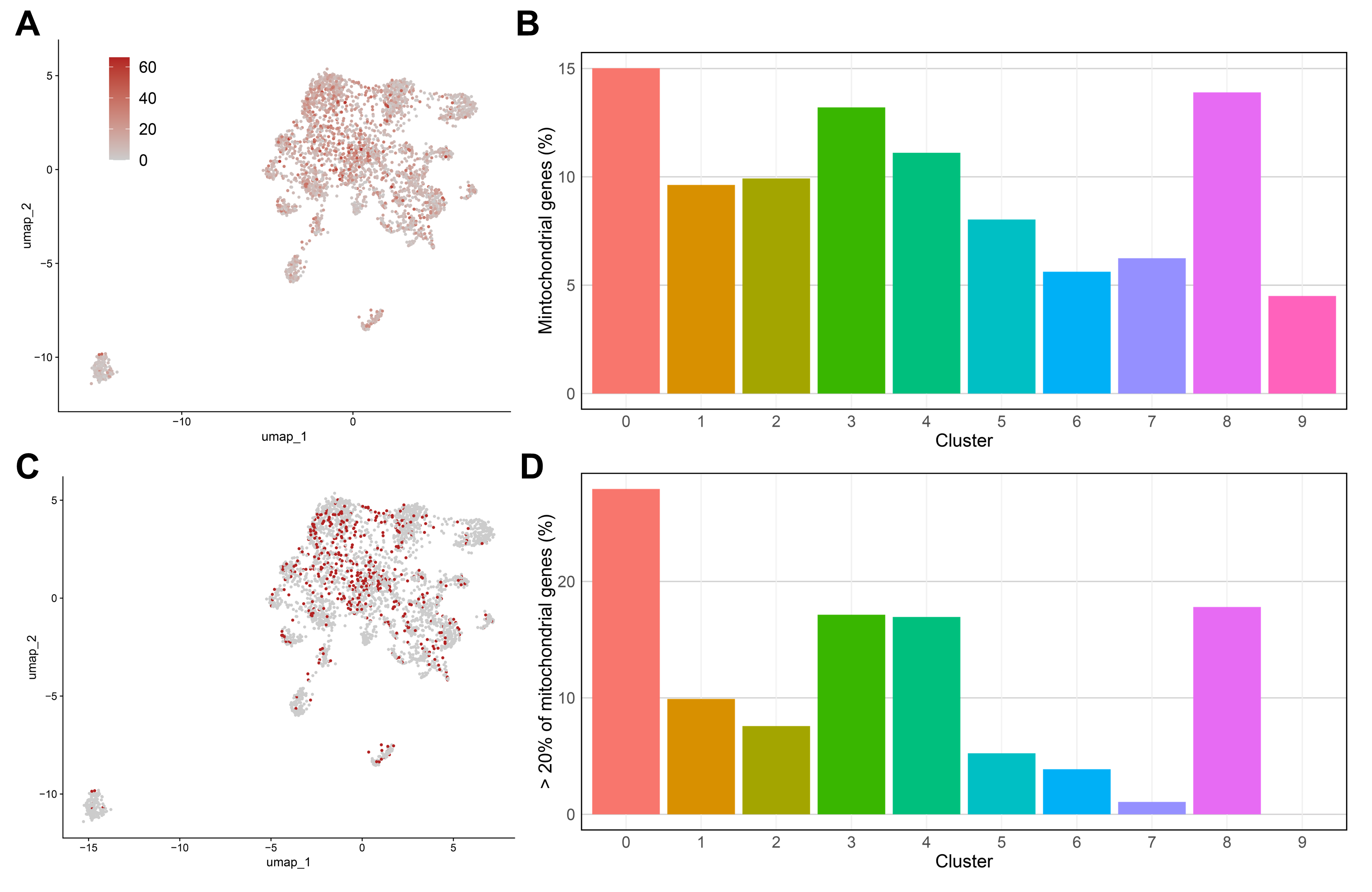

### Fig. S3

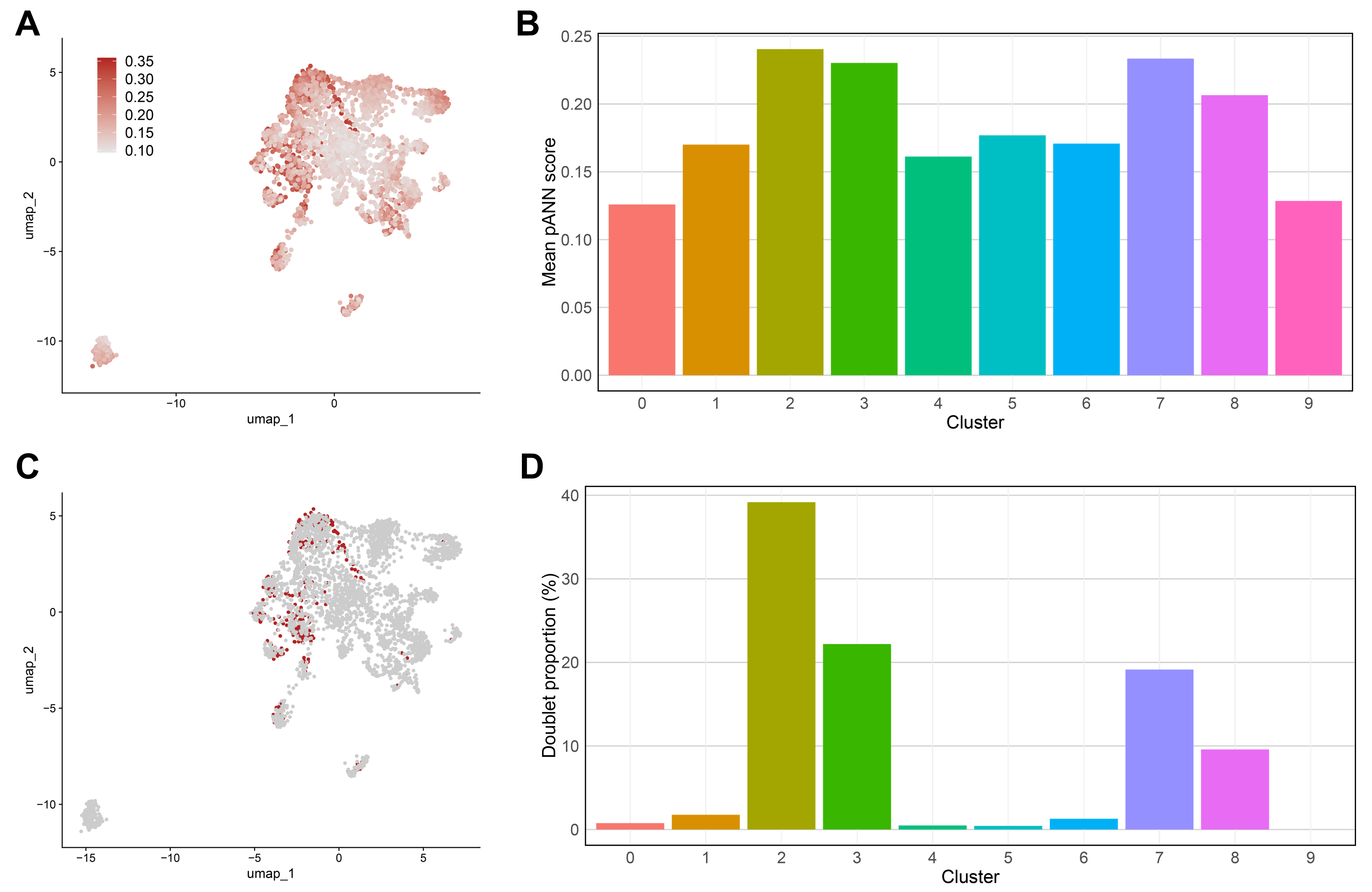

### Fig. S4

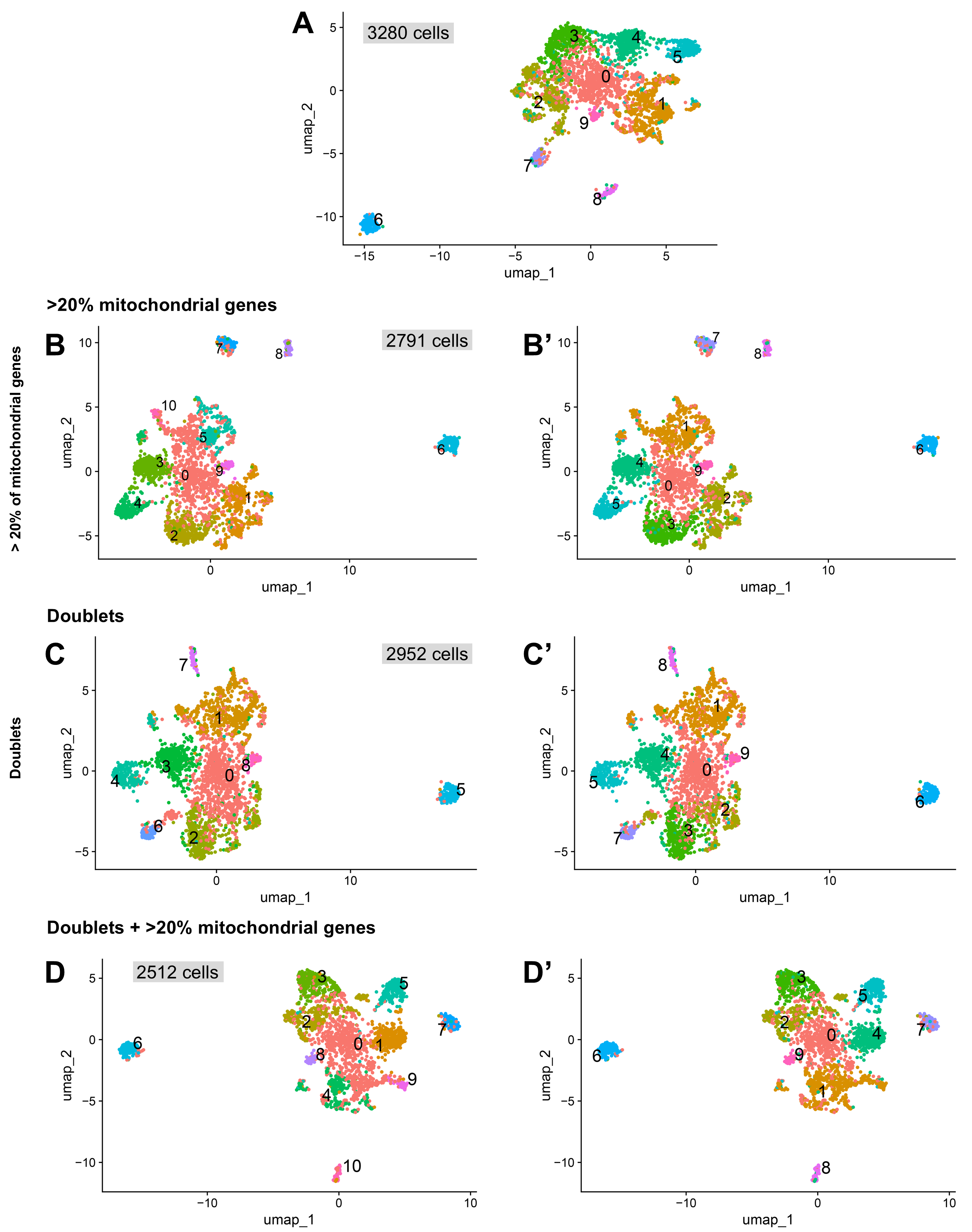
